# Peptidoglycan DD-peptidases have distinct activities that impact fitness of *Acinetobacter baumannii*

**DOI:** 10.1101/2025.07.25.666485

**Authors:** Arshya F. Tehrani, Abhisha Khadka, Berenice Furlan, Christine Pybus, Michael Whalen, Jacob Biboy, Orietta Massidda, Waldemar Vollmer, Joseph M. Boll

## Abstract

The Gram-negative cell envelope is a vital interface between the bacterial cell and its environment. It acts as a selective barrier, blocking harmful agents while permitting nutrient uptake. Additionally, it enables environmental sensing and adaptive responses. Structurally, it is composed of the outer membrane, the cytoplasmic (inner) membrane, and the periplasm, which contains the peptidoglycan layer. Peptidoglycan is a conserved polymer that provides structural integrity, allowing the cell to withstand the internal turgor. It consists of glycan strands connected by short peptides, forming a mesh-like structure. In Gram-negative bacteria, the majority of the peptidoglycan subunits contain tetrapeptides. Tetrapeptides are generated through the action of DD-carboxypeptidases (DD-CPases), which cleave the terminal D-alanine from pentapeptides. Gram-negative bacteria encode multiple DD-CPases, but their precise role in maintaining cell shape and structural integrity remain poorly understood. The nosocomial pathogen *Acinetobacter baumannii* encodes three putative DD-CPases. To investigate the role of these enzymes, we generated single mutants, as well as double mutants in *dacC*, *dacD,* and *pbpG,* which encode the homologs of *Escherichia coli* DD-CPases PBP5, PBP6a, PBP6b, and the endopeptidase (DD-EPase) PBP7, respectively. We assessed the mutants for changes in cell morphology, growth dynamics, and stress tolerance. Additionally, we analyzed the composition of their peptidoglycan layers to determine the biochemical consequences of their inactivation. Each mutant exhibited distinct alterations in coccobacillary morphology and growth. Peptidoglycan analysis confirmed the enzymes possess DD-CPase activity, and PBP6b also demonstrated endopeptidase activity. Together, our results demonstrate that each peptidoglycan-modifying enzyme contributes uniquely to cell growth and morphology. These findings underscore their non-redundant functions and suggest their specific activities may serve as valuable targets for developing new antimicrobial therapies.

**Importance:** DD-peptidases, including carboxypeptidases and endopeptidases are crucial for maintaining cell envelope homeostasis, with distinct roles for each enzyme in cell wall biogenesis and structural integrity. The enzymatic characterization presented in this study not only advance our understanding of fundamental *A. baumannii* biology but also highlight these enzymatic activities as targets for development of innovative therapeutic strategies to combat infections caused by this multidrug-resistant microbe.

## Introduction

The Gram-negative cell envelope is structurally organized into three distinct layers, each critical for maintaining cellular integrity and environmental fitness. The hallmark of Gram-negative bacteria is the asymmetric outer membrane (OM), which consists of an inner leaflet of glycerophospholipids, and an outer (surface-exposed) leaflet enriched with lipopolysaccharide (LPS) or lipooligosaccharide (LOS). The lipid asymmetry forms a robust permeability barrier that restricts the entry of toxic compounds (1). Beneath the OM lies the cytoplasmic (inner) membrane, a symmetric bilayer composed of glycerophospholipids. Between the two membranes is the periplasmic space, which contains a thin peptidoglycan (PG) layer (2). The PG is the primary determinant of cell shape and mechanical strength, protecting the cell from osmotic lysis. However, recent studies suggest that the LPS-rich OM also contributes significantly to the mechanical stability and shape maintenance during osmotic changes (3, 4). Despite these insights, the molecular mechanisms that coordinate the biogenesis and integration of the OM and PG layers remain poorly understood. Emerging evidence points to direct interactions between PG maturation and β-barrel assembly machinery (BAM) complex, suggesting a physical and functional linkage connecting the envelope layers (5). Disrupting these connections may represent a promising strategy to compromise envelope integrity and sensitize bacteria to environmental and antibiotic stress.

Penicillin-binding proteins (PBPs) are a diverse group of enzymes essential for the biosynthesis, remodeling, and maintenance of the PG layer. These enzymes catalyze key reactions such as formation or breaking of peptide bonds, which are critical for forming and modifying peptide cross-links within the PG meshwork. PBPs are broadly categorized into high molecular weight (HMW) and low molecular weight (LMW) groups, each with distinct enzymatic functions that contribute to PG assembly and homeostasis (6). HMW PBPs are PG synthases and further divided into two classes: Class A PBPs, which possess both glycosyltransferase and transpeptidase activities, and Class B PBP, which function as transpeptidases (6, 7). The former enzymes are responsible for polymerizing glycan strands and cross-linking peptides, thereby constructing the PG scaffold. LMW PBPs are generally non-essential under standard growth conditions but play important roles in PG maturation, remodeling, and turnover (8–12). They function mainly as DD-endopeptidases (DD-EPases) and/or DD-carboxypeptidases (DD-CPases) cleaving amide bonds within stem peptides to regulate PG architecture and facilitate cell wall plasticity during growth and environmental adaptation (13–15).

DD-CPases are critical enzymes involved in the maturation and remodeling of the PG layer. These enzymes hydrolyze the peptide bond between the fourth and fifth amino acids (both usually D-Ala) of pentapeptides, thereby releasing a terminal D-alanine and generating tetrapeptides (16). This PG maturation occurs within minutes of the incorporation of new PG strands in *E.coli* (17). Tetrapeptides, in turn, serve as substrates for various downstream cell envelope processes. In *Acinetobacter baumannii*, for instance, tetrapeptides are utilized by the LD-transpeptidase (LD-TPase) LdtJ to form alternative 3-3 crosslinks and add D-amino acids to position four of tetrapeptides in PG (18, 19). Beyond their role in PG architecture, tetrapeptides have been shown to bind BAM and inhibit the assembly of OM proteins, dampening OM protein assembly near mature PG (e.g., the old poles) during the cell cycle (5). Moreover, tetrapeptide formation has been implicated in establishing physical and functional connections between the PG layer and OM proteins, contributing to overall stability of the cell envelope via LD-TPase-mediated attachment of the OM-anchored lipoprotein Lpp in *E. coli*. DD-CPases are broadly conserved across Gram-negative bacteria (20), with orthologs identified in Enterobacterales, *Pseudomonas aeruginosa*, and *A. baumannii*.

*E. coli* encodes at least seven DD-CPases, including PBP4 (also has DD-EPase activity), PBP4b, PBP5, PBP6a, PBP6b, AmpC and AmpH (21–25), as summarized in Table S1. In addition, class A PBPs can also have considerable DD-CPase activity (26). PBP5, encoded by *dacA*, is the most abundant DD-CPase in *E. coli* (27). Deletion of *dacA* along with at least two other DD-CPase genes results in minor morphological abnormalities, such as kinks, bends, or branches (28). However, significant morphological defects only arise when all DD-CPase genes are deleted, in addition to the deletion of *mrcA* (encoding PBP1A), highlighting their overlapping yet distinct functions in PG maintenance. When *E. coli* grows at low pH conditions, PBP6b takes over as the most important DD-CPase from PBP5 (29). Under conditions of severe outer membrane biogenesis stress the seemingly ‘minor’ PBP6b, but not PBP5, becomes essential (30). The weak beta-lactamase AmpC and the beta-lactamase AmpH also contribute to shape maintenance. Whether the other DD-CPases have specialized functions is not known.

In contrast, *A. baumannii* encodes only three predicted DD-CPase genes. *dacC* encodes a protein with high homology to both *E. coli* PBP5 (42% identity) and PBP6a (41%identitiy), denoted herein as PBP5/6a, *dacD* encodes PBP6b, and *pbpG* encodes PBP7, with the latter having both DD-EPase and DD-CPase activity (31), as summarized in Table S1. The AmpC beta-lactamase (encoded by *bla_ADC-26_*) also exhibits minor DD-CPase activity (32). While *A. baumannii* PBP7 has both DD-EPase and DD-CPase activity (31), *E. coli* PBP7 has only DD-endopeptidase activity and has a role in cell division (33, 34). In *A. baumannii* PBP7 has been shown to be important for bacterial survival during antibiotic treatment, although the precise mechanism underlying the increased susceptibility upon its inactivation remains unclear (31). The functional redundancy in *E. coli* contrasts with the streamlined DD-CPase system in *A. baumannii*, making it a powerful model for dissecting the specific contributions of individual DD-CPases to cell envelope integrity and bacterial fitness.

In this study, we generated targeted mutations in genes encoding the predicted DD-CPases, including *dacC* and *dacD*, the predicted DD-CPase/DD-EPase *pbpG*, and the predicted beta-lactamase *bla_ADC-26_*to investigate their individual and combined roles. We systematically assessed the morphological and physiological consequences of single and double mutations by analyzing changes in cell shape, growth kinetics, and sensitivity to environmental stress. Additionally, we analyzed the PG composition in the single mutants and biochemically defined the enzymatic activities of PBP5/6a and PBP6b. Our findings demonstrate that each DD-CPase contributes uniquely to *A. baumannii* growth, morphology, and overall fitness. These enzymes function in distinct, non-redundant pathways to support PG biosynthesis and homeostasis across different phases of the cell cycle.

## Results

### Morphological defects associated with deletion of *dacC*, *dacD*, or *pbpG*

Given that *A. baumannii* encodes only three-four predicted DD-CPases, relative to at least seven in *E. coli* (29), we hypothesized that single gene deletions in *A. baumannii* may result in distinct morphological phenotypes, offering fundamental insights into the specific roles of these enzymes.

Morphological defects were evident in each *A. baumannii* DD-CPase mutant (Figure 1). The Δ*dacC* mutant exhibited a population of short, spherical cells relative to wild-type strain ATCC 17978 (Figure 1A). Quantitative measurements confirmed a significant reduction in cell length and width (Figure 1B; Figure S1A), accompanied by slower growth (Figure S1B). Complementation of PBP5/6a from a non-native locus rescued the cell morphology and restored normal growth (Figure 1A, 1B, S1A, and S1B).

**Figure 1:**
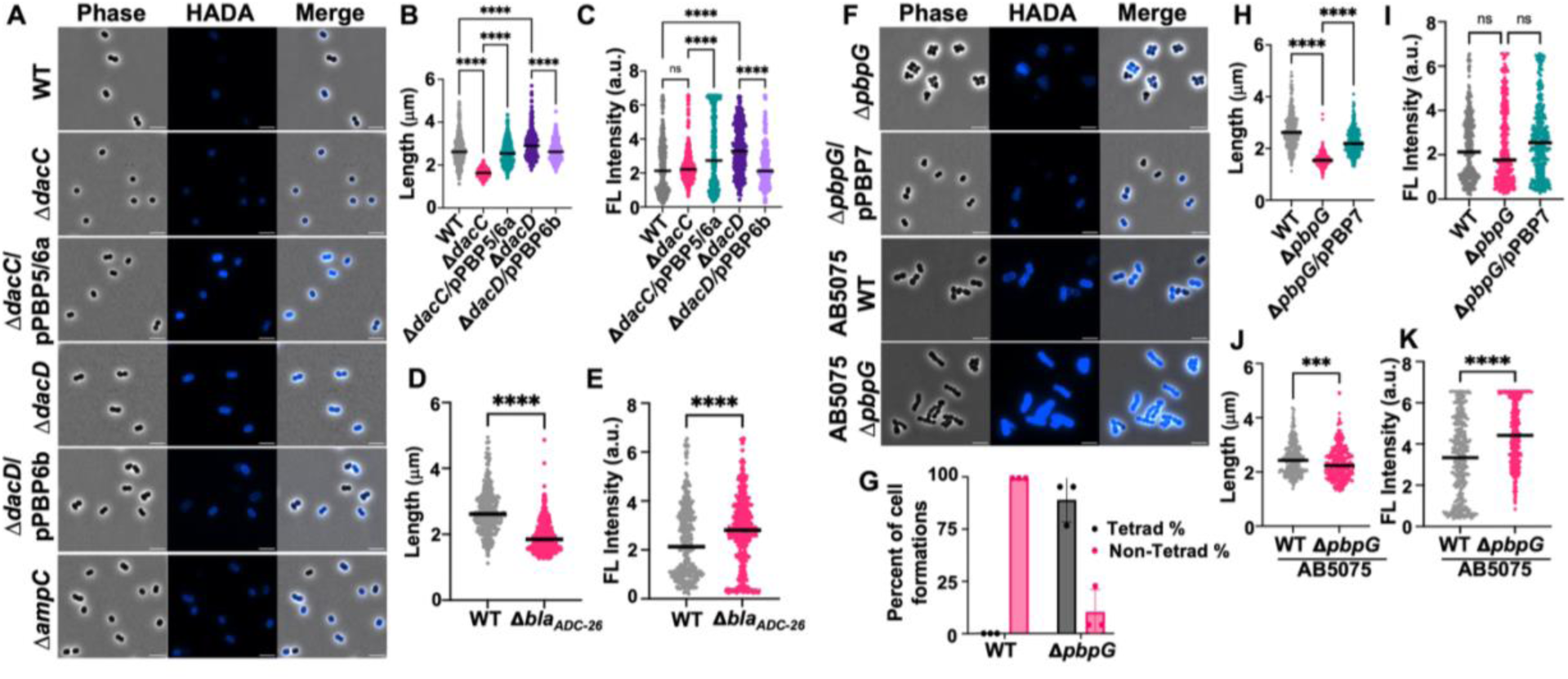
Microscopy of *A. baumannii* mutants in logarithmic growth phase. **(A)** Phase-contrast (left), fluorescence (middle), and merged (right) images of wild-type (WT) strain ATCC 17978, Δ*dacC*, Δ*dacC*/pPBP5/6a, Δ*dacD*, Δ*dacD*/pPBP6b, and Δ*bla_ADC-26_* cells. Scale bar: 5 μm. (**B)** Quantification of cell length (pole to pole) quantifications of each cell population (*n* ≥ 300), measured using ImageJ software with MicrobeJ plugin. Each dot represents a single cell. Error bars indicate standard deviation. Statistical significance was determined using one-way ANOVA (*** *P* < 0.001, **** *P* < 0.0001). **(C)** Fluorescence intensity quantifications for Δ*dacC* and Δ*dacD***. (D)** Quantification of cell length for Δ*bla_ADC-26_*, as described in (B). **(E)** Fluorescence intensity quantifications for Δ*bla_ADC-26_***. (F)** Microscopy images, as described in (A), showing strains ATCC 17978 Δ*pbpG,* Δ*pbpG*/pPBP7, AB5075 WT, and AB5075 Δ*pbpG*. **(G)** Quantification of coccobacilli and tetrad formations in strain ATCC 17978 WT and Δ*pbpG* using ImageJ software (*n* ≥300). Tetrads are reported as a percentage of the total population. Data were collected from three independent experiments; one representative image and dataset are shown. **(H)** Quantification of cell length for strain ATCC 17978 Δ*pbpG*, as described in (B). **(I)** Fluorescence intensity quantifications for strain ATCC 17978 Δ*pbpG***. (J)** Quantification of cell length for strain AB5075 WT and Δ*pbpG*, as described in (B). **(K)** Fluorescence intensity quantifications for strain AB5075 WT *and* Δ*pbpG*.

In contrast, the Δ*dacD* mutant displayed an elongated cell morphology with some heterogeneity (Figure 1A and 1B; Figure S1A), and increased D-amino acid modification of PG (Figure 1C). Cells were stained with a fluorescent derivative of D-alanine (HADA) (35), which is incorporated into PG by PBPs and LD-TPases (36–39), and enables D-amino acid modification measurements. Increased modification with D-alanine may arise from increased tetrapeptide pools which are substrates for the LD-TPase LdtJ (18) — or from loss of PBP6b-mediated D-amino acid trimming from pentaapeptides. While the precise mechanism remains unclear, these findings suggest a regulatory role for PBP6b in D-amino acid incorporation into position five of the peptide. Growth rate was like wild-type (Figure S1C), and complementation fully restored the wild-type morphology and fluorescence pattern (Figure 1A–C; S1A).

We also examined the role of the AmpC β-lactamase, (Acinetobacter-derived cephalosporinase (ADC), encoded by *bla_ADC-26_*), which has been reported to exhibit weak DD-CPase activity (32). Δ*bla_ADC-26_*mutants displayed variable morphologies, with enrichment of spherical cells (Figure 1A, 1D, and S1D), resembling the Δ*dacC* phenotype (Figure 1B and S1A), but with greater variability in cell length. Whole-genome sequencing of several Δ*bla_ADC-26_* isolates revealed suppressor mutations, none of which were associated with PG structural components known to influence cell shape. Due to this genetic heterogeneity, the Δ*bla_ADC-26_* mutant was excluded from further analysis.

To assess strain-specific effects, we analyzed DD-CPase mutants in a second *A. baumannii* background, the well characterized clinical isolate AB5075 (40), using transposon mutants from the Manoil library (41). Similar shape and growth defects were observed in the AB5075 Δ*dacC* and Δ*dacD* mutants (Figure S2), supporting the same effects for enzymatic inactivation across different strains, with the exception that the Δ*dacD* mutant did not show an elongated cell morphology in strain AB5075. Length, width, fluorescence intensity, and growth curve calculations were similar to trends described in strain ATCC 17978 (Figure S2A, B, C, D, and E).

Our previous work demonstrated that PBP7 (encoded by *pbpG*) exhibits DD-CPase and DD-EPase activity and is important for antibiotic survival (31). In strain ATCC 17978, Δ*pbpG* mutants formed spherical cells that clustered into tetrads, suggesting altered septum positioning and/or impaired cell separation (Figure 1F, 1G). These cells also showed reduced length (Figure 1H), width (Figure S3A), and slightly decreased fluorescence upon treatment with HADA (Figure 1I). Complementation restored wild-type morphology and growth (Figure 1F, 1H, and 1I; S3A-B). This phenotype may reflect a combined lack of DD-CPase and DD-EPase activities.

In strain AB5075, Δ*pbpG* mutants also formed spherical cells but did not form tetrads (Figure 1F). Instead, they formed chains of rounder cells (Figure 1J and S3C), indicating a defect in cell elongation and separation. Cells also showed increased HADA fluorescence (Figure 1K), suggesting increased PG D-amino acid modification and/or reduced removal. The strain-specific differences suggest that while PBP7 plays a consistent enzymatic role affecting morphology, additional functions may vary in different *A. baumannii* genetic backgrounds.

### Morphological and growth defects associated with combined deletions of *dacC*, *dacD*, and *pbpG*

While single DD-CPase mutants in *A. baumannii* exhibit distinct morphological and growth phenotypes, previous studies in *E. coli* have shown that only multiple deletions (28), particularly in the Δ*dacA* background, are required to elicit significant morphological defects, suggesting functional redundancy among DD-CPases. To determine whether paired deletions in *A. baumannii* would exacerbate phenotypic defects, we engineered the Δ*dacC* Δ*dacD*, Δ*dacC* Δ*pbpG*, and Δ*dacD* Δ*pbpG* double mutants in strain ATCC 17978 (Figure 2). Surprisingly, the double mutants did not exhibit exacerbated phenotypes. Instead, each double mutant was characterized by a combination of the respective single mutant characteristics. In the Δ*dacC* Δ*dacD* mutant (Figure 2A), cells appeared spherical, like Δ*dacC* cells, but with increased morphological heterogeneity (Figure 2B) and interestingly, the additional deletion of *dacD* rescued the slower growth observed in the Δ*dacC* single mutant (Figure S3D). HADA incorporation was significantly higher relative to wild type (Figure 2C), resembling the Δ*dacD* phenotype (Figure 1C). These findings suggest that PBP5/6a and PBP6b have distinct functions: PBP5/6a in cell elongation, whereas PBP6b in affecting D-amino acid incorporation into PG. The increase in D-amino acid modification may compensate for the loss of PBP5/6a, thereby restoring normal growth.

**Figure 2:**
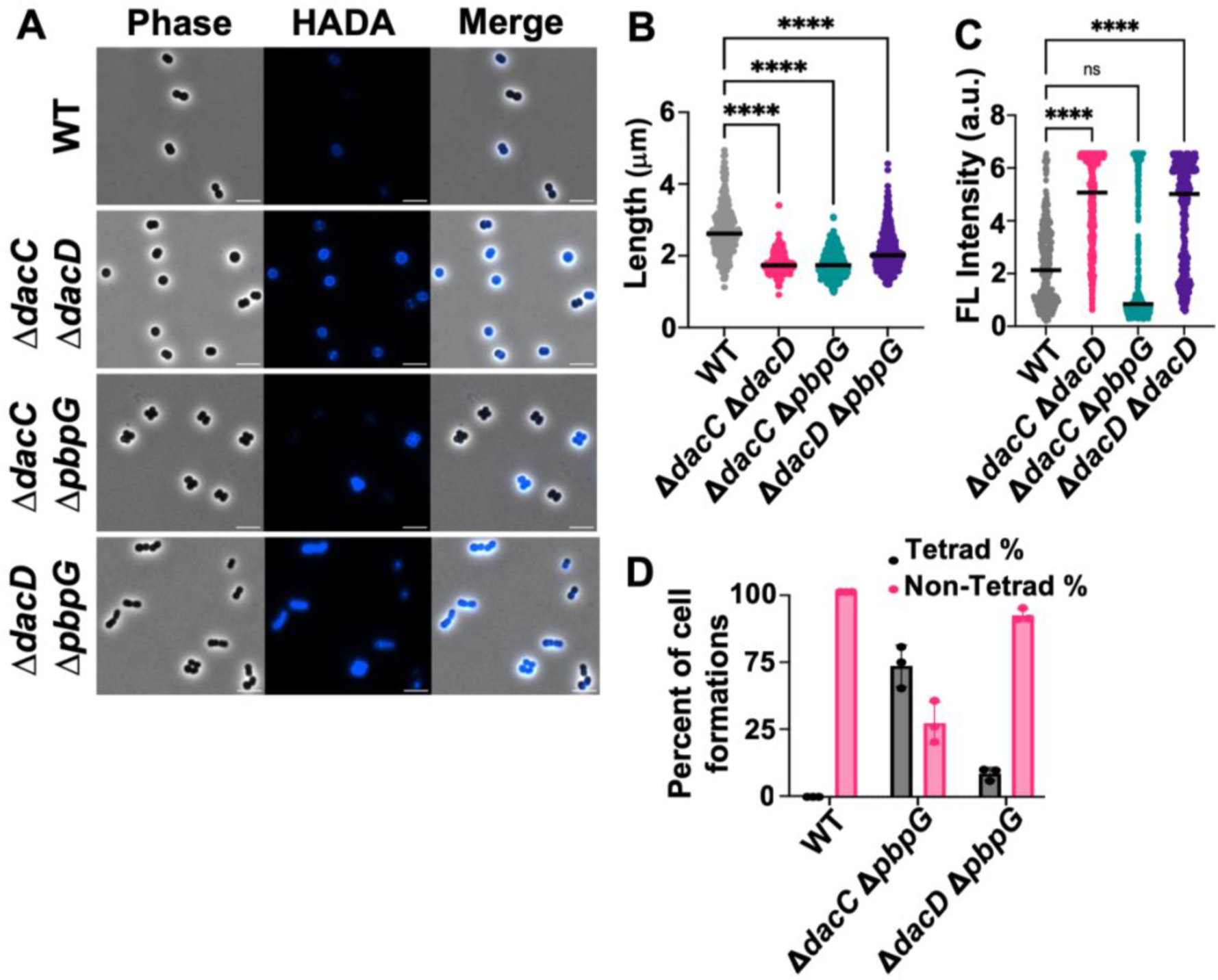
Microscopy of *A. baumannii* strain 17978 double mutants. **(A)** Phase-contrast (left) fluorescence (middle) and merged (right) images of wild-type (WT), Δ*dacC* Δ*dacD*, Δ*dacC* Δ*pbpG*, Δ*dacD* Δ*pbpG*. Scale bar: 5 μm. **(B)** Quantification of cell length (pole to pole) quantifications of each cell population (*n* ≥ 300) was calculated using ImageJ software with MicrobeJ plugin. Each dot represents one cell. Error bars represent standard deviation. Statistical significance was determined using one-way ANOVA (**** *P* < 0.0001). **C)** Quantification of fluorescence intensity. **(D)** Quantification of coccobacilli and tetrad cell formations using ImageJ software (*n* ≥ 300) Tetrads are reported as a percentage of the total population. Data were collected from three independent experiments; one representative image and dataset are shown.

In the Δ*dacC* Δ*pbpG* double mutants, the cells were spherical (Figure 2A and 2B), consistent with elongation defects observed in both single mutants (Figure 1A). However, the phenotype was not more severe, implying that PBP5/6a and PBP7 may have overlapping roles in cell elongation. HADA fluorescence intensity remained similar to wild type (Figure 2C). Notably, tetrad cell clusters, indicative of altered septum positioning and/or cell separation defects, were formed (Figure 2A) mirroring the Δ*pbpG* phenotype (Figure 1F and 1K). A growth defect was also observed, and after 12 h, cell growth appeared compromised (Figure S3D), suggesting a synthetic fitness defect in the double mutant.

The Δ*dacD* Δ*pbpG* mutant displayed short, spherical cells similar to the Δ*pbpG* single mutant cells (Figure 2A and 2B). However, unlike in other Δ*pbpG* mutants in the ATCC 17978 background, these cells did not consistently form tetrads. While some tetrads were present, most cells existed as single short units or chains (Figure 2D), indicating that Δ*dacD* disruption partially rescues the defect associated with *pbpG* deletion. This mutant also showed improved growth (Figure S2D) and elevated HADA fluorescence intensity, consistent with increased D-amino acid incorporation or less removal (Figure 2C). Together, these results suggest that *A. baumannii* DD-CPases have both distinct and overlapping roles in PG biosynthesis and remodeling and envelope homeostasis. The absence of additive morphological defects in DD-CPase double mutants highlights the complexity of their functions and suggests compensatory mechanisms that buffer against loss of individual enzymes.

### Muropeptide analysis in the DD-CPase mutants reveals altered PG structures

To investigate how PBP5/6a and PBP6b contribute to *A. baumannii* PG structure, we analyzed the muropeptide compositions of wild type, Δ*dacC,* and Δ*dacD* mutants in exponential and stationary phases (Figure 3, Table S2). PG was isolated and digested with a muramidase, and the resulting muropeptides were reduced with sodium borohydride and analyzed by high-performance liquid chromatography (HPLC), as previously described (18, 31, 42). Each mutant exhibited distinct muropeptide profiles relative to wild type. Wild type strain ATCC 17978 showed the typical profile of a tetrapeptide-rich PG with only minor differences between growth phases (Figure 3A and 3D). In contrast, Δ*dacC* mutants displayed the presence of more pentapeptide-containing muropeptides during both exponential and stationary phases (Figure 3B and 3E), indicating that PBP5/6a is required for trimming D-alanine residues from pentapeptides – a hallmark of DD-CPase activity (16).

**Figure 3:**
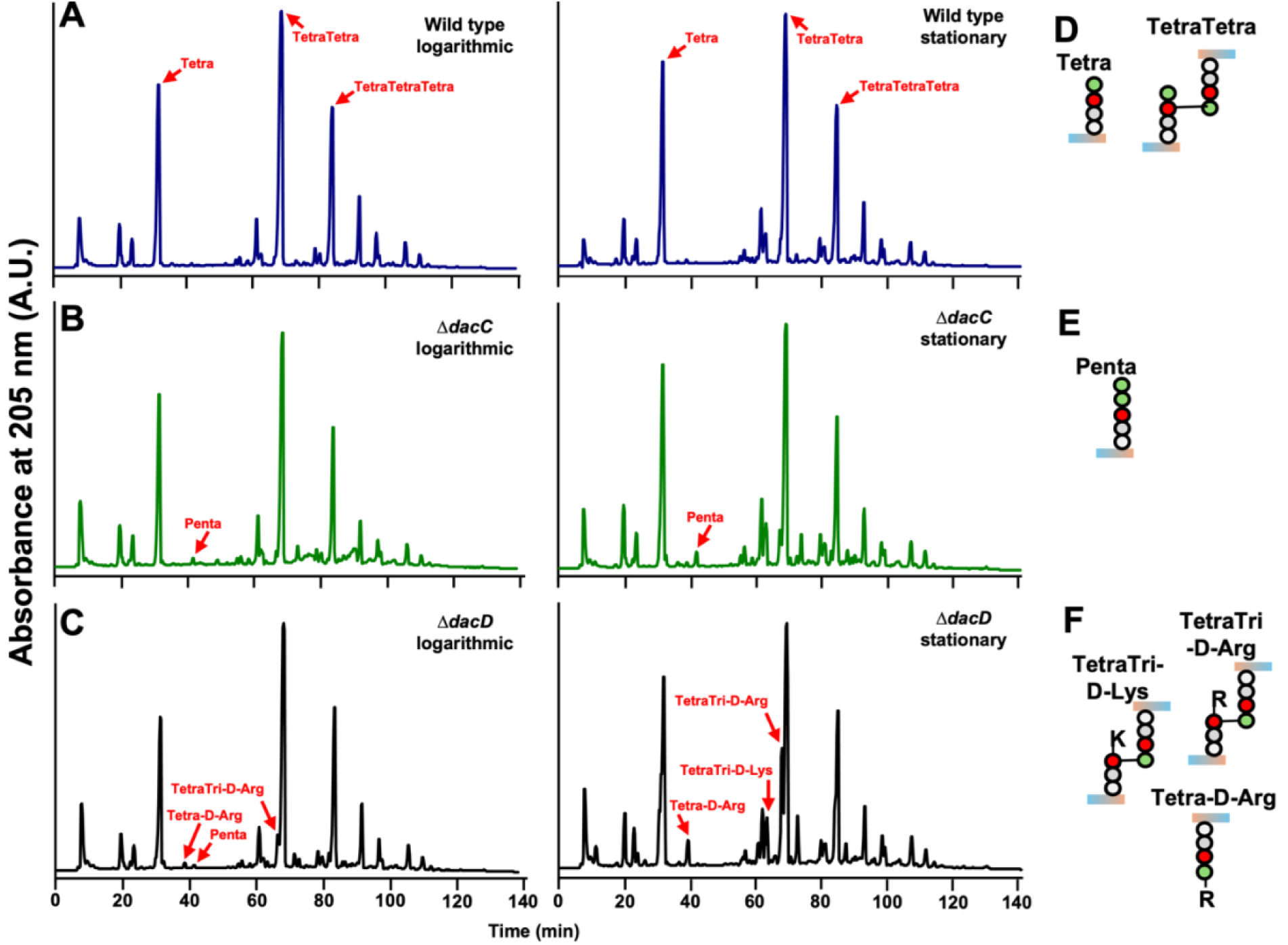
Muropeptide analysis in *A. baumannii* strain 17978 Δ*dacC* and Δ*dacD* strains. Peptidoglycan was isolated from **(A)** wild type, **(B)** Δ*dacC*, and **(C)** Δ*dacD* strains during logarithmic (left) or stationary (right) growth phases, digested with muramidase and the resulting muropeptides analyzed by high-performance liquid chromatography (HPLC). The Δ*dacC* mutant exhibited enrichment of pentapeptides, while the Δ*dacD* mutant showed increased levels of D-amino acid modified muropeptides relative to wild type. Red arrows indicate peaks corresponding to specific muropeptide structures. **(D, E, and F)** Structural illustration of the labeled muropeptides shown in panels (A-C).

Δ*dacD* mutants also showed pentapeptide enrichment during exponential growth, consistent with DD-CPase activity. However, in both growth phases, these mutants exhibited elevated levels of D-amino acid-modified tetrapeptides (Figure 3C and 3F). This increase correlates with the enhanced fluorescence observed in Δ*dacD* cells labeled HADA (Figure 1A and 1C), suggesting that PBP6b may regulate D-amino acids incorporation into PG.

Peptides with D-amino acids (e.g., HADA) at position four and five are formed by exchange reactions with the terminal D-Ala on tetrapeptides and pentapeptides, respectively. PBPs exchange D-amino acids at position five resulting in tetrapeptide-D-amino acid (tetrapeptides with an additional amino acid at position five), LD-TPases exchange D-amino acids at position four, resulting in tripeptide-D-amino acids. DD-CPases are only expected to remove D-amino acids from position five. Hence, D-amino acid-modified tetrapeptides could be enriched due to loss of PBP6b-mediated trimming of D-amino acids from pentapeptides. Notably, pentapeptide enrichment was only observed during exponential growth, suggesting that PBP6b functions as a DD-CPase primarily during active cell wall synthesis, while promoting D-amino acid removal during stationary phase.

Together, these results support distinct roles for PBP5/6a and PBP6b in PG remodeling: PBP5/6a primarily trims pentapeptides to tetrapeptides and regulates cell elongation, while PBP6b modulates D-amino acid incorporation, in the stationary phase, both potentially influencing PG cross-linking and envelope stability.

### PBP5/6a and PBP6b are DD-CPases that catalyze tetrapeptide formation, and PBP6b is also an EPase

To validate the proposed enzymatic functions of PBP5/6a and PBP6b supported by PG analysis, we performed activity assays using the purified recombinant enzymes (Figure S4). Native PBP5/6a and PBP6b, along with their respective catalytically inactive mutants (PBP5/6a_S121A_ and PBP6b_S122A_), in which the catalytic active serine was substituted with alanine, were expressed and purified. Each enzyme was incubated with muropeptides from *E. coli* CS703-1 - enriched in pentapeptides - as previously described (31), and reactions were conducted at both pH 7.5 and pH 5.0 to assess pH-dependent activity.

PBP5/6a exhibited robust DD-CPase activity as evidenced by a marked reduction in pentapeptide peaks and a corresponding increase in tetrapeptides at both pH values (Figure 4A). No activity was observed in the reactions containing the PBP5/6a_S121A_ active site mutant or in the no-enzyme controls. Additionally, PBP5/6a reduced TetraPenta levels while increasing TetraTetra muropeptides, further supporting its DD-CPase activity.

**Figure 4:**
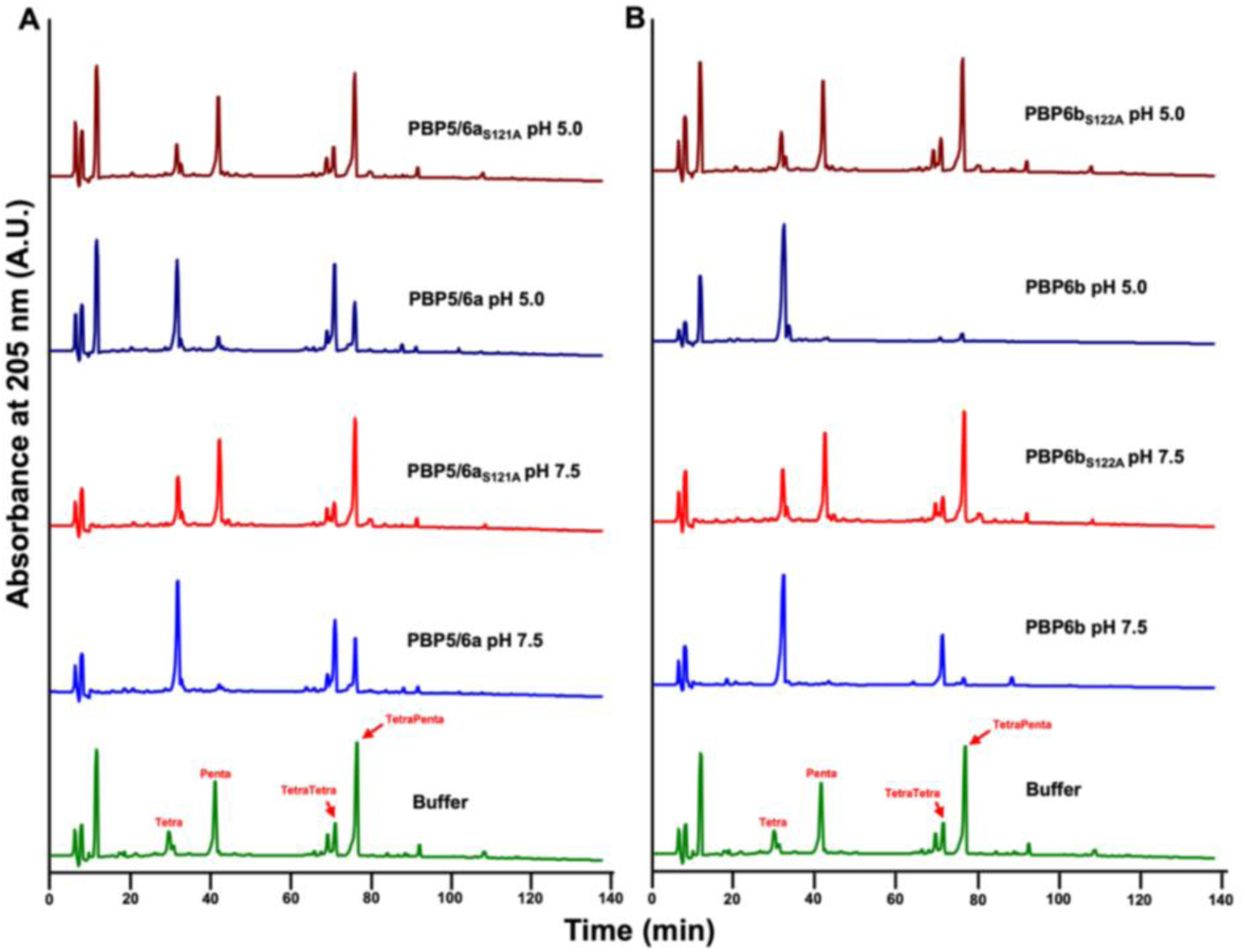
PBP5/6a and PBP6b exhibit DD-carboxypeptidase activity, and PBP6b also functions as an endopeptidase. **(A)** Recombinant PBP5/6a and its active site mutant PBP5/6a_S121A_, and **(B)** PBP6b and its active site mutant PBP6b_S122A_, were incubated with peptidoglycan isolated from *E. coli* D456 (CS703-1), which contains Tetra, Penta, TetraTetra, and TetraPenta as the major components. Both PBP5/6a and PBP6b exhibited DD-carboxypeptidase activity, cleaving pentapeptides, whereas the active-site mutants showed no activity. PBP6b also demonstrated endopeptidase activity at both neutral (pH 7.5) and acidic pH (pH 5.0) conditions. Enzymes were used at a final concentration of 10 μM. Muropeptides are labeled in red.

PBP6b was also active against pentapeptides and tetrapentapeptides converting these to Tetra and TetraTetra-muropeptides, respectively, at pH 7.5 (Figure 4B). Interestingly, under acidic conditions at pH 5.0 more so than at pH 7.5, PBP6b showed robust DD-EPase activity. This dual activity was dependent on the catalytic serine residue, as the PBP6b_S122A_ active site mutant and no-enzyme controls showed no such changes.

Together, these results confirm that PBP5/6a and PBP6b catalyze formation of tetrapeptides from pentapeptides, consistent with their role as DD-CPases anticipated by sequence homology and PG analysis. Moreover, PBP6b exhibits pH-dependent DD-EPase activity, potentially contributing to PG remodeling under acidic conditions, as previously observed for *E. coli* PBP6b, which only showed DD-CPase activity (29).

### PBP5/6a and PBP7 play unique roles under pH stress

To determine whether *A. baumannii* DD-CPases contribute to environmental stress tolerance, we assessed the growth and cell morphology defects of Δ*dacC,* Δ*dacD*, Δ*pbpG* mutants under acidic or alkaline conditions (Figure 5). The Δ*dacC* mutant exhibited growth defects in acidic (pH 5.0) and alkaline conditions (pH 9.2), showing reduced growth in both relative to neutral pH (pH 7.5) (Figure 5A). These findings suggest that PBP5/6a is required for maintaining cell envelope integrity across a broad pH range, and that its function cannot be compensated by other DD-CPases.

**Figure 5:**
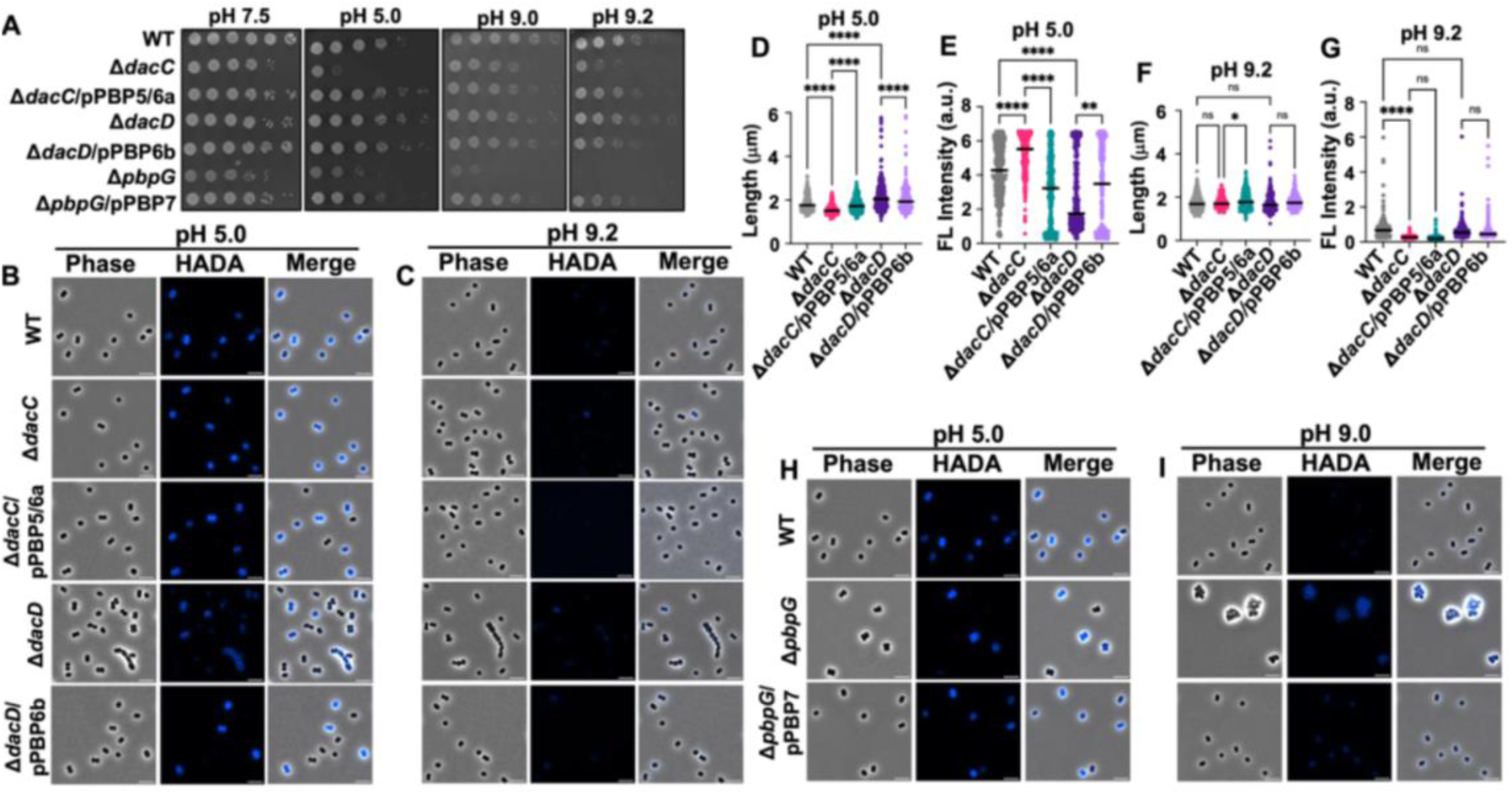
Growth and morphology of *A. baumannii* strain ATCC 17978 mutants under acidic and alkaline stress conditions. **(A)** Colony spot assays of wild-type (WT) and mutant strains serially diluted 10-fold starting at OD_600_ 0.05, plated on LB agar adjusted to the indicated pH values. **(B)** Phase-contrast (left), fluorescence (middle), and merged images of WT, Δ*dacC*, Δ*dacC*/pPBP5/6a, Δ*dacD*, and Δ*dacD*/pPBP6b cells grown at pH 5.0. **(C)** Same as (B), but cells grown at pH 9.2. Scale bar: 5 μm. **(D)** Quantification of cell length (pole to pole) at pH 5.0 of cell populations (*n* ≥300), measured using ImageJ with the MicrobeJ plugin. Each dot represents a single cell. Error bars indicate standard deviation. Statistical significance was determined using one-way ANOVA (* *P* < 0.05, ** *P* <0.01, **** *P* <0.0001). **(E)** Quantification of fluorescence intensity at pH 5.0. **(F)** Same as (D), but for cells grown at pH 9.0. **(G)** Same as (E), but for cells grown at pH 9.0. **(H)** Phase-contrast (left,) fluorescence (middle), and merged (right) images of WT, Δ*pbpG* and Δ*pbpG*/pPBP7 cells grown at pH 5.0. **(I)** Same as (H), but cells grown at pH 9.0. Each experiment was independently replicated three times, and one representative data set was reported.

In contrast, the Δ*dacD* mutant did not show obvious growth defects under any tested condition (Figure 5A). Interestingly, this mutant displayed enhanced tolerance to alkaline and acidic pH relative to wild-type, suggesting that PBP6b inactivation confers a selective advantage in stress environments, potentially by altering PG remodeling or D-amino acid incorporation (Figure 3C and 4B).

The Δ*pbpG* mutant was highly sensitive to both acidic and alkaline conditions, with complete growth inhibition at pH 9.2 (Figure 5A). This indicates that PBP7 is essential for survival under pH stress and, like PBP5/6a, its function is not redundant with other DD-CPases.

### DD-Carboxypeptidase mutants exhibit unique morphological responses under pH stress

To further investigate the role of DD-CPases under environmental stress, we examined the morphology of Δ*dacC,* Δ*dacD*, Δ*pbpG* cells using microscopy under acidic (Figure 5B, 5D, 5H, & S5A) or alkaline (Figure 5C, 5F, 5G, 5I, & S5B) conditions. In alkaline conditions, all strains – including wild-type – exhibited enrichment of spherical cells (Figure S5C, S5D, S5F, S5H, S5I, & S5K) and reduced HADA fluorescence intensity (Figure S5E, S5G, & S5J) compared to neutral pH. These observations suggest that alkaline stress may affect cell elongation and D-amino acid modification of PG, likely through mechanisms independent of PBP5/6a and PBP6b. In contrast, acidic conditions led to shorter, brighter cells in the ATCC 17978 wild-type strain, consistent with reduced D-amino acid removal (Figure S5E, S5G, & S5J). Mutants displayed similar length phenotypes at acidic pH relative to those observed at a neutral pH: Δ*dacC* cells were shorter, and Δ*dacD* cells were longer (Figure 5D). However, a notable difference was the occasional appearance of chaining phenotypes in Δ*dacD* cells under both pH extremes (Figure 5B, 5C, & 5D), suggesting a potential role for PBP6b in cell separation during stress. HADA fluorescence intensity analysis revealed a striking shift at pH 5.0: Δ*dacC* cells show increased fluorescence intensity relative to wild type, while Δ*dacD* cells showed reduced fluorescence (Figure 5E). This suggests that PBP5/6a may remove incorporated D-amino acids under acidic conditions, a role previously attributed to PBP6b at neutral pH.

In the Δ*pbpG* mutants, tetrad-like cell clusters persisted under both acidic and alkaline conditions (Figure 5H and 5I), but the nature of these clusters varied. Under acidic conditions, tetrads appeared as a single group of clustered cells, indicating a severe defect. In alkaline conditions multiple tetrads aggregated into larger clusters, further compounding the phenotype. These morphological abnormalities likely contribute to the complete growth inhibition at pH 9.2 (Figure 5A). The distinct responses under each stress conditions suggest that PBP7 may have additional, condition-specific roles in regulating cell wall synthesis and envelope remodeling.

## Discussion

PG homeostasis - through coordinated synthesis, remodeling and recycling - is essential for bacterial survival, morphology, and adaptation to environmental stress. Among the enzymes involved in PG remodeling and turnover, DD-CPases remain to be further studied in Gram-negative bacteria other than *E. coli*. These enzymes trim terminal D-alanine residues from pentapeptides, generating tetrapeptides that influence PG cross-linking, cell shape, and stress resilience. In *E. coli*, DD-CPases are functionally redundant, with multiple enzymes compensating for one another depending on environmental conditions (29).

In this study, we characterized the roles of the putative DD-CPases PBP5/6a and PBP6b, and the DD-CPase/EPase PBP7 in *A. baumannii*, revealing different roles than their homologs in *E. coli*. Our results demonstrate that each DD-CPase in *A. baumannii* performs a unique, non-redundant function in PG homeostasis, cell shape maintenance, and environmental stress adaptation.

The Δ*dacC* mutant exhibited short spherical cells, implicating PBP5/6a in cell elongation. Δ*dacD* mutants, while slightly longer than wild-type, displayed increased D-amino acid incorporation, suggesting that PBP6b may remove D-amino acids added by PBP-mediated exchange. Δ*pbpG* mutants exhibited the most severe defects, forming tetrad-like clusters, consistent with an important role for PBP7 in cell morphology and separation. These observed phenotypes were consistent also in the clinical isolate AB5075, although with some strain-specific morphological variation in the Δ*dacD* and, particularly, the Δ*pbpG* mutants.

Double mutant analysis further supported the non-redundant nature of these enzymes. Combinatorial deletions (Δ*dacC* Δ*dacD*, Δ*dacC* Δ*pbpG*, Δ*dacD* Δ*pbpG*) resulted in additive or compensatory phenotypes, but not in synthetic lethality. Notably, Δ*dacD* Δ*pbpG* mutants showed partial restoration of the Δ*pbpG* phenotype, suggesting that PBP6b may compensate for the morphological defects associated with PBP7 loss.

Muropeptide profiling revealed that Δ*dacC* mutants accumulated pentapeptides, confirming PBP5/6a’s role in pentapeptide trimming. Δ*dacD* mutants showed enrichment of D-arginine and D-lysine acid-modified muropeptides, suggesting that PBP6b may specifically cleaves these modified substrates or indirectly regulate their incorporation. These findings highlight the specialized substrate preferences and regulatory roles of individual DD-CPases in *A. baumannii*.

Enzymatic assays confirmed that both PBP5/6a and PBP6b possess DD-CPase activity. Additionally, PBP6b exhibited DD-EPase activity that increased at acidic pH, similar to the previously reported DD-EPase activity for PBP7 (31). This reduced specificity—combining DD-CPase and DD-EPase activities—may explain the lack of redundancy among *A. baumannii* DD-CPases, in contrast to the environment-specific redundancy observed in *E. coli* aside from PBP4’s DD-CPase/EPase activity (29).

Environmental stress assays further emphasized the unique roles of these enzymes. Δ*dacC* and Δ*pbpG* mutants exhibited growth defects across all tested pH conditions, while Δ*dacD* mutants showed enhanced tolerance to alkaline stress. Morphological analysis under pH stress revealed that Δ*pbpG* mutants experienced exacerbated shape defects, with tetrads forming as single unseparated cells under acidic conditions and as aggregated clusters under alkaline conditions. These observations suggest that PBP7 may have additional, condition-specific roles in cell wall remodeling.

In conclusion, our study reveals that *A. baumannii* DD-peptidases are not functionally redundant but instead perform distinct, crucial roles in PG remodeling, cell shape maintenance, and stress adaptation. The multifunctionality of these enzymes—combining DD-CPase and DD-EPase activities—may reflect an evolutionary adaptation to streamline PG regulation in this clinically relevant pathogen. Future studies should explore the broader regulatory networks and potential therapeutic vulnerabilities associated with these unique enzymes.

## Materials and Methods

### Bacterial strains and growth conditions

Strains and plasmids used in this study are listed in Table S3. *Acinetobacter baumannii* strains were cultured aerobically from frozen stock on Luria-Bertani (LB) agar or in LB broth at 37°C. Unless otherwise specified, antibiotics were used at the following concentrations: 25 mg/L kanamycin, and 10 mg/L tetracycline.

*A. baumannii* strain ATCC 17978 has been reported to consist of two distinct variants (43). To confirm the variant used in this study, colony PCR was performed using primers specific to the cardiolipin synthase gene (*clsC2*), which yielded a positive product, indicating that all experiments were conducted using the *A. baumannii* 17978 UN variant. *A. baumannii* strains, including AB5075, are known to exhibit phase-variable colony opacity phenotypes (44). Microscopic examination of AB5075 cultures revealed that most colonies were opaque, with only <1% translucent colonies observed among more that 200 colonies per plate (*n* = 3 plates per strain). For consistency, only opaque variants were used in all experiments, including those involving the AB5075 *dacA/C::tn*, *pbp6b::tn*, and *pbpG::tn* mutants.

### Construction of genetic mutants

The primers used in this study are listed in Table S4 *A. baumannii* mutants, including Δ*dacC*, Δ*dacD*, Δ*pbpG*, Δ*bla_ADC-26_*, and all double mutants were generated using the recombination-mediated genetic engineering (recombineering), as previously described (18, 45–47). Briefly, a kanamycin resistance cassette flanked by FLP recombination target (FRT) sites was PCR-amplified from the pKD4 plasmid using primers containing 125-bp homology arms specific to the target gene. The linear PCR product was electroporated into *A. baumannii* strain ATCC 17978, harboring the pRECAb plasmid (pAT03). Following recovery in LB broth, transformants were plated on LB agar supplemented with 7.5 mg/L kanamycin. All mutants were verified by PCR.

To remove the recombineering plasmid, mutants were grown on LB agar containing 2 mM nickel(II) chloride (NiCl₂) and replica-plated onto LB agar with either kanamycin or tetracycline, as previously done (42). Colonies that were kanamycin-resistant and tetracycline-sensitive were confirmed by PCR to have lost the plasmid.

For markerless deletions, the kanamycin resistance cassette was excised by transforming the cured mutants with pMMB67EH::FLP (pAT08), which expresses FLP recombinase. Transformants were plated on LB agar containing tetracycline and 2 mM IPTG to induce recombinase expression. Successful excision of the resistance cassette was confirmed by PCR.

For complementation, the coding sequences of *dacC* (PBP5/6a), *dacD* (PBP6b), and *pbpG* (PBP7), including 200 bp of upstream and downstream flanking regions, were amplified from *A. baumannii* ATCC 17978 genomic DNA. These fragments were cloned into the pABBRknR plasmid at the XhoI and KpnI restriction sites. The resulting constructs (pPBP5/6a, pPBP6b, and pPBP7) were introduced into their respective deletion mutants to restore gene expression under the control of the native promoter.

### Fluorescent HADA staining

FDAA staining was performed as previously described (18, 31, 48). Overnight cultures were grown with shaking at 37°C in 5 mL of LB broth, supplemented with appropriate antibiotics when necessary. The following day, cultures were diluted 1:100 into fresh LB medium (5 mL total volume) and incubated at 37°C with shaking until OD_600_ 0.3-0.5. The cells were harvested, washed to remove residual debris, and resuspended in 1 mL LB. Two microliters of 10 mM HCC- (linezolid-7-nitrobenz-2-oxa-1,3-diazol-4-yl)-amino-d-alanine (HADA) were added to each culture. The mixtures were incubated at 37°C for 20 minutes to allow incorporation of the fluorescent alanine. Following staining, cultures were washed with LB broth and fixed in phosphate-buffered saline (PBS) containing a 1:10 dilution of 16% paraformaldehyde.

### Microscopy

Microscopy was performed as previously described (18, 31, 48). Paraformaldehyde-fixed cells were mounted on 1.5% agarose pads and imaged using an inverted Nikon Eclipse Ti-2 wide-field epifluorescence microscope. The system was equipped with a Photometrics Prime 95B camera and a Plan Apo 100× objective lens (NA 1.45). Phase-contrast and fluorescence images were acquired using NIS Elements software. Blue fluorescence was visualized using a Sola LED light engine with the following filter set: 350/50nm excitation filter, 460/50nm emission filter, and a 400nm dichroic mirror.

### Image analysis

Microscopy images were processed as previously described (18, 31, 48). Pseudocoloring of fluorescence images was performed using ImageJ Fiji (49). Quantitative measurements, including length, width, fluorescence intensity, and total cell surface area, were measured using MicrobeJ (18, 31, 48, 50). Data were analyzed and visualized using GraphPad Prism version 10.2.2. Each experiment was independently repeated three times. Quantification data and representative images shown in the figures are from one representative biological replicate.

### Peptidoglycan isolation

PG was isolated as previously described (18, 31, 42). Biological replicates were cultured in 400 mL of LB broth to either mid-logarithmic or stationary phase. Cells were harvested by centrifugation at 7,000 x *g* for 20 minutes at 4°C using an Avanti JXN-26 Beckman Coulter centrifuge with a JA-10 rotor. Pellets were resuspended in 6 mL of chilled PBS and lysed by dropwise addition of boiling 8% sodium dodecyl sulfate (SDS). PG was purified following the protocol established by Glauner (51). Sacculi were treated with the Cellosyl muramidase (gift from Hoechst, Frankfurt am Main, Germany), to release muropeptides, which were then reduced with sodium borohydride and separated using high-performance liquid chromatography (HPLC) on a 250 × 4.6 mm 3-μm Prontosil 120-3-C_18_ AQ reversed-phase column (Bischoff, Leonberg, Germany). Elution was monitored by absorbance at 205 nm. Muropeptide peaks were identified by comparison to published chromatograms (18, 42, 52).

### Construction of PBP5/6a and PBP6b active-site mutants

To generate catalytically inactive variants of PBP5/6a and PBP6b, gene fragments encoding PBP5/6a_S121A_ (serine 121 substituted with alanine) and PBP6b_S122A_ (serine 122 substituted with alanine) were synthesized by Twist Biosciences. These mutations target the conserved serine residue in the active site, essential for enzymatic activity.

### Construction of PBP5/6a and PBP6b overexpression strains

The coding sequences for *dacC* and *dacD* were amplified from *A. baumannii* ATCC 17978 genomic DNA, while *dacC*_S121A_ and *dacD*_S122A_ were amplified from synthesized gene fragment DNA. The amplicons were generated using primers that included a C-terminal His_8×_-tag sequence to facilitate downstream purification. The resulting PCR products were cloned into the NdeI and BamHI restriction sites of the pT7-7Kn expression vector (53). The constructs pT7-7Kn::*dacC*, pT7-7Kn::*dacC*_S121A_, pT7-7Kn::*dacD*, and pT7-7Kn::*dacD*_S122A_ were transformed into *E. coli* C2987 chemically competent cells and confirmed by Sanger sequencing. Verified plasmids were subsequently transformed into *E. coli* C2527 (BL21) cells (New England BioLabs, Inc.) for protein expression and purification.

### Purification of recombinant PBP5/6a and PBP6b

PBP expression and purification was done as previously described (31). Briefly, *E. coli* BL21 cells harboring pT7-7Kn::*dacC*, pT7-7Kn::*dacC*_S121A_ and pT7-7Kn::*dacD*_S122A_ were cultured in 1000 mL of LB broth at 37°C until reaching OD_600_ 0.3-0.5. Due to toxicity associated with overexpression, 4000 mL of culture was needed for *E. coli* BL21 cells harboring pT7-7Kn::*dacD*. 1L was sufficient with the other cultures. All cultures were induced with 1 mM IPTG for 4 hours at 25°C.

Following incubation, cells were harvested, washed with cold 1× PBS, and pelleted. Pellets were frozen overnight at −20°C, then thawed on ice and resuspended in 20 mL of lysis buffer (20 mM Tris, 300 mM NaCl, 10 mM imidazole, pH 8). Cells were lysed by sonication using a Fisher Scientific Model 120 Sonic Dismembrator, with 20 seconds on, followed by 20 seconds off, for a total of 10 cycles at 80% amplitude. Lysates were centrifuged at 15,428 x *g* for 10 minutes at 4°C. Pellets were resuspended in extraction buffer (25mM Tris/HCl, 10mM MgCl_2_ 1M NaCl, 0.02% NaN_3_, 2% TX-100, pH 7.5) and incubated overnight at 4°C with gentle rocking. The next day, supernatants were collected with centrifugation and incubated with HisPur Ni-nitrilotriacetic acid (NTA) resin (Thermo Scientific), pre-equilibrated with extraction buffer for 2 h at 4°C with rotation. The resin mixture was transferred to a 10-mL gravity-flow column and washed sequentially with 20 mL of lysis buffer containing increasing imidazole concentrations (0 mM, 15 mM, and 30 mM). Proteins were eluted in 8 fractions using 500 µL of elution buffer (20 mM Tris, 300 mM NaCl, 250 mM imidazole, pH 8), incubated for 5 minutes before gravity elution. Fractions containing protein, identified by SDS-PAGE, were pooled and dialyzed overnight at 4°C in dialysis buffer (10 mM Tris, 50 mM KCl, 0.1 mM EDTA, 5% glycerol, pH 8) using 12-mL dialysis cassettes (Thermo Scientific). Protein purity and identity were confirmed by SDS-PAGE and bocillin-binding assays.

### Activity assays

PBP5/6a and PBP6b activity assays were performed in a final reaction volume of 50 μL containing either: Neutral pH buffer: 20mM Tris/HCl, 50mM NaCl, 2mM MgCl_2_ pH 7.5, or acidic pH buffer: 20mM sodium acetate, 50mM NaCl, 2mM MgCl_2,_ pH 5.0. Each reaction contained 10 μM of purified PBP5/6a, PBP5/6a_S121A_, PBP6b, or PBP6b_S122A_. PG isolated from *E. coli* CS703-1, enriched in pentapeptides was added as a substrate. Reactions were incubated at 37°C for 16 hours. Samples were boiled for 10 minutes to terminate reactions and incubated with 100 μg/mL cellosyl for further 16 hours at 37°C. Samples were then boiled for 10 minutes and centrifuged at 16,000xg for 10 minutes in a microfuge. The resulting muropeptides were reduced with sodium borohydride and acidified to a pH of 4.0–4.5. Control samples contained PG from *E. coli* CS703-1 without enzyme. Muropeptide analysis was performed as previously described (51).

### Spot assays with pH medium

Overnight cultures were grown in LB broth at 37°C and then backdiluted to OD 0.05. Cultures were serially diluted 1:10 across five dilution steps. From each dilution, 5 uL aliquots were spotted onto LB agar plates adjusted to various pH levels. To prepare pH-adjusted LB plates, 50mM Tris was added to standard LB agar, and the pH was adjusted to the desired value using either HCl or NaOH. Plates were incubated at 37°C for 16 h, after which images were captured. All images shown are representative of three independent experiments.

### Statistical Analysis

Statistical analysis for cell morphology and fluorescence intensity data was performed using one-way ANOVA. A *P* value of less that 0.05 was considered statistically significant.

## Acknowledgements

The work was supported by funding from the National Institutes of Health (grants R35GM143053 to J.M.B. and R01AI168159 to J.M.B. and W.V.) and the UK BBSRC (BB/W013630/1 to W.V.). We thank Dr. Daniela Vollmer for the preparation of peptidoglycan.

---

**Figure S1:**
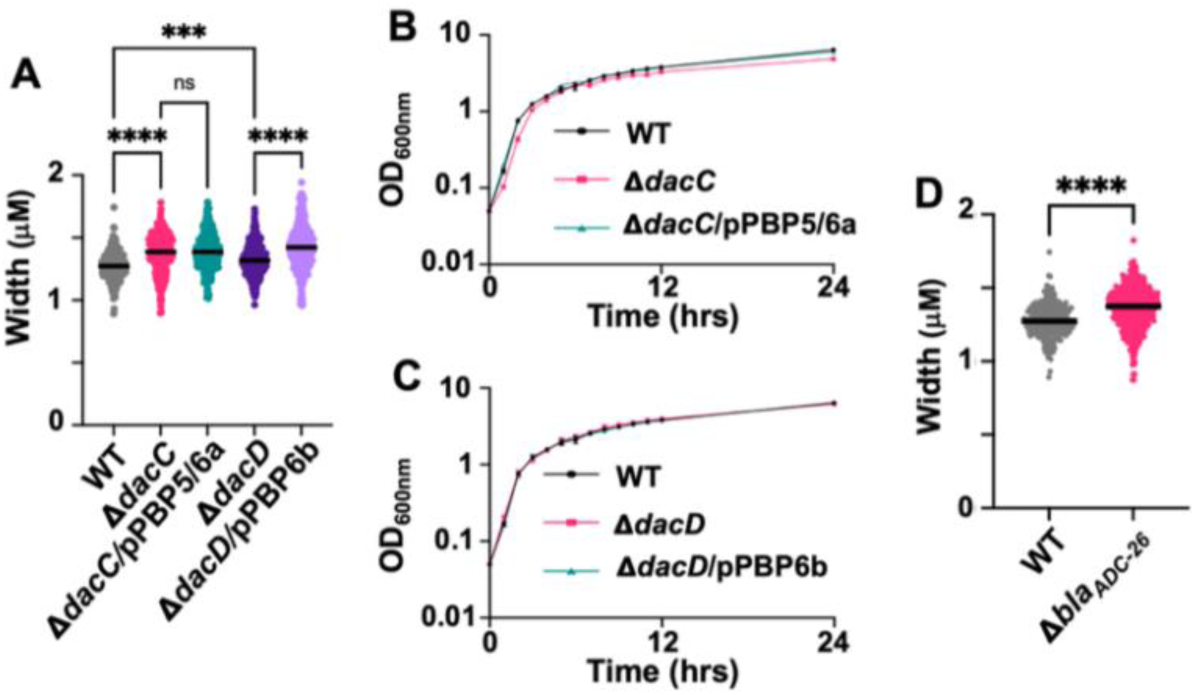
Analysis of Δ*dacC* and Δ*dacD* mutations in *A. baumannii* strain ATCC 17978. **(A)** Quantifications of cell width in wild type (WT), Δ*dacC*, Δ*dacC*/pPBP5/6a, Δ*dacD*, Δ*dacD*/pPBP6b and Δ*bla_ADC-26_* strains (*n* ≥300), measured using ImageJ with the MicrobeJ plugin. Each dot represents a single cell. Error bars indicate standard deviation. Statistical significance was determined using one-way ANOVA (*** *P* < 0.001, **** *P* < 0.0001). **(B)** Growth curves of the Δ*dacC* mutant. **(C)** Growth curves of the Δ*dacD* mutant. (**D)** Quantification of cell width in the Δ*bla_ADC-26_* mutant, as described in (A).

**Figure S2:**
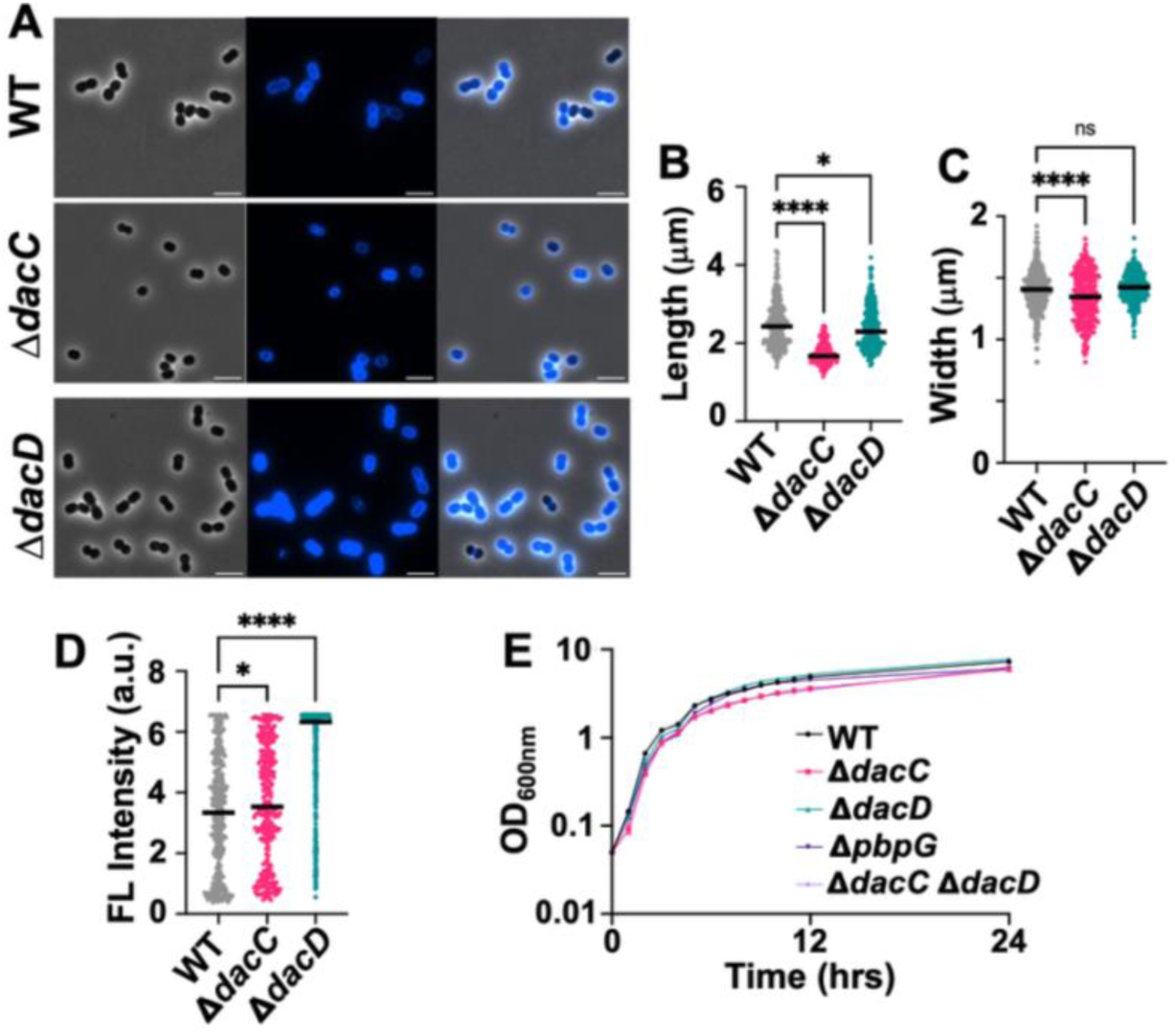
Analysis of *A. baumannii* strain AB5075 mutants. **(A)** Phase-contrast (left) fluorescence (middle) and merged (right) images of wild-type (WT), Δ*dacC*, Δ*dacD* cells. Scale bar: 5 μm. (**B)** Quantification of cell length (pole to pole) for each strain (*n* ≥300) was measured using ImageJ with the MicrobeJ plugin. Each dot represents a single cell. Error bars indicate standard deviation. Statistical significance was determined using one-way ANOVA (* *P* < 0.05, **** *P* < 0.0001). **(C)** Quantification of cell width. **(D)** Quantification of fluorescence intensity. **(E)** Growth curves of Δ*dacC,* Δ*dacD,* and Δ*pbpG* mutants. Each experiment was independently replicated three times; one representative data set is shown.

**Figure S3:**
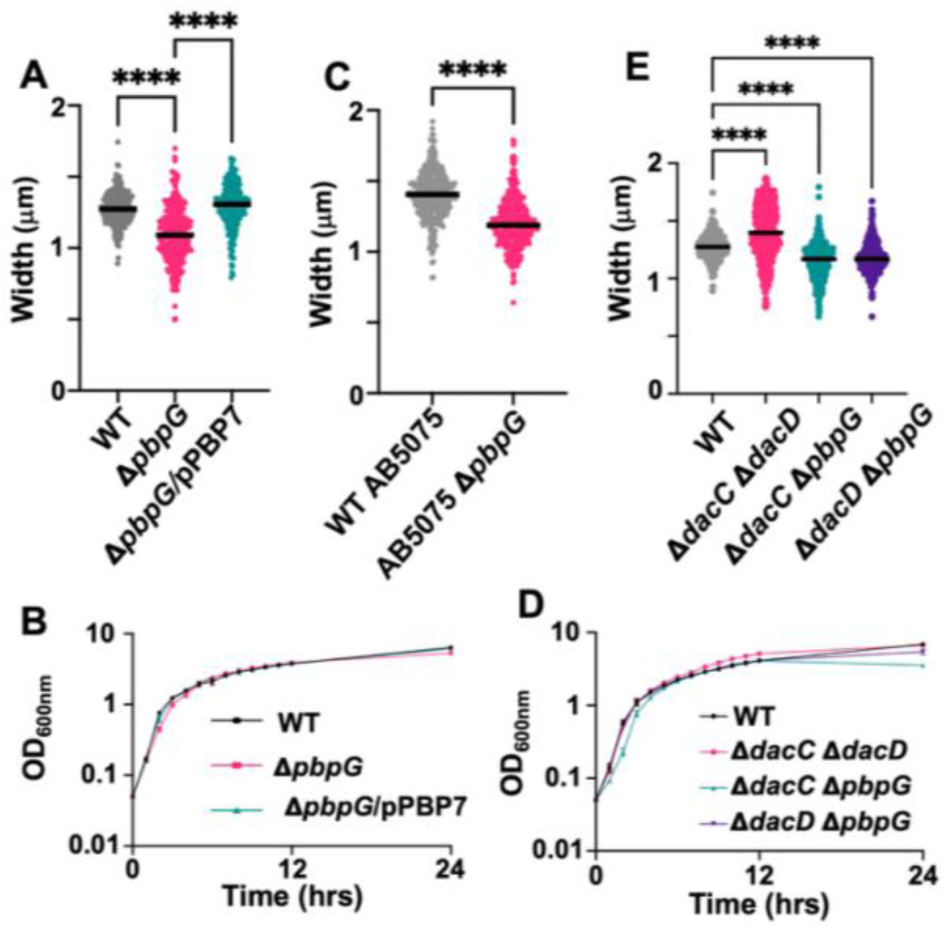
Cell morphology measurement and growth curves of Δ*pbpG* mutants and double mutants. **(A)** Quantifications of cell width in *A. baumannii* strain ATCC 17978 Δ*pbpG* mutants (*n* ≥300), measured using ImageJ with the MicrobeJ plugin. Each dot represents a single cell. Error bars represent standard deviation. Statistical significance was determined using one-way ANOVA (**** *P* < 0.0001). **(B)** Growth curves of Δ*pbpG* mutants in strain ATCC 17978. **(C)** Same as described in (A), but for Δ*pbpG* mutants in *A. baumannii* strain AB5075. **(D)** Growth curves of double mutants in strain *A. baumannii* ATCC 17978. **(E)** Same as described in (A), but for double mutants in *A. baumannii* strain ATCC 17978.

**Figure S4:**
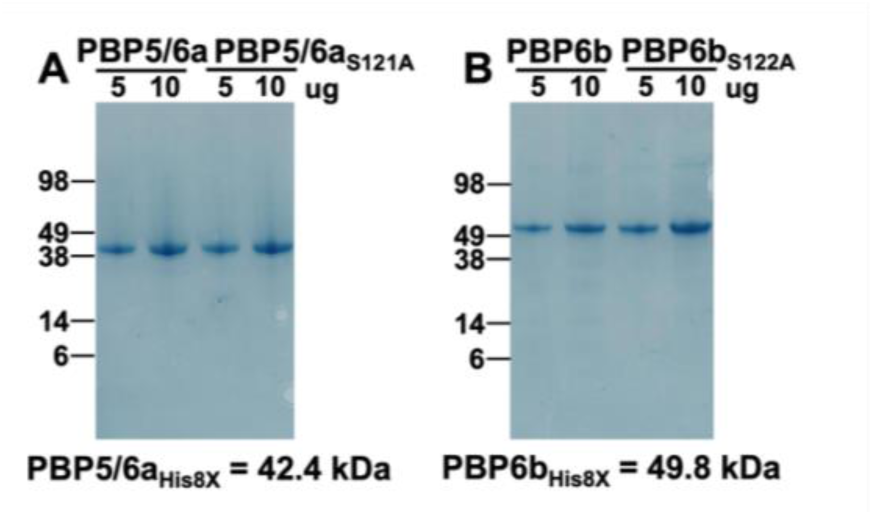
Purification of PBP5/6a and PBP6b enzymes. **(A)** Coomassie stained SDS-PAGE gel of PBP5/6a_His8X_ and the catalytically inactive mutant PBP5/6a_S121A His8X_. **(B)** Coomassie stained SDS-PAGE gel of PBP6b_His8X_ and the catalytically inactive mutant PBP6b_S122A His8X_.

**Figure S5:**
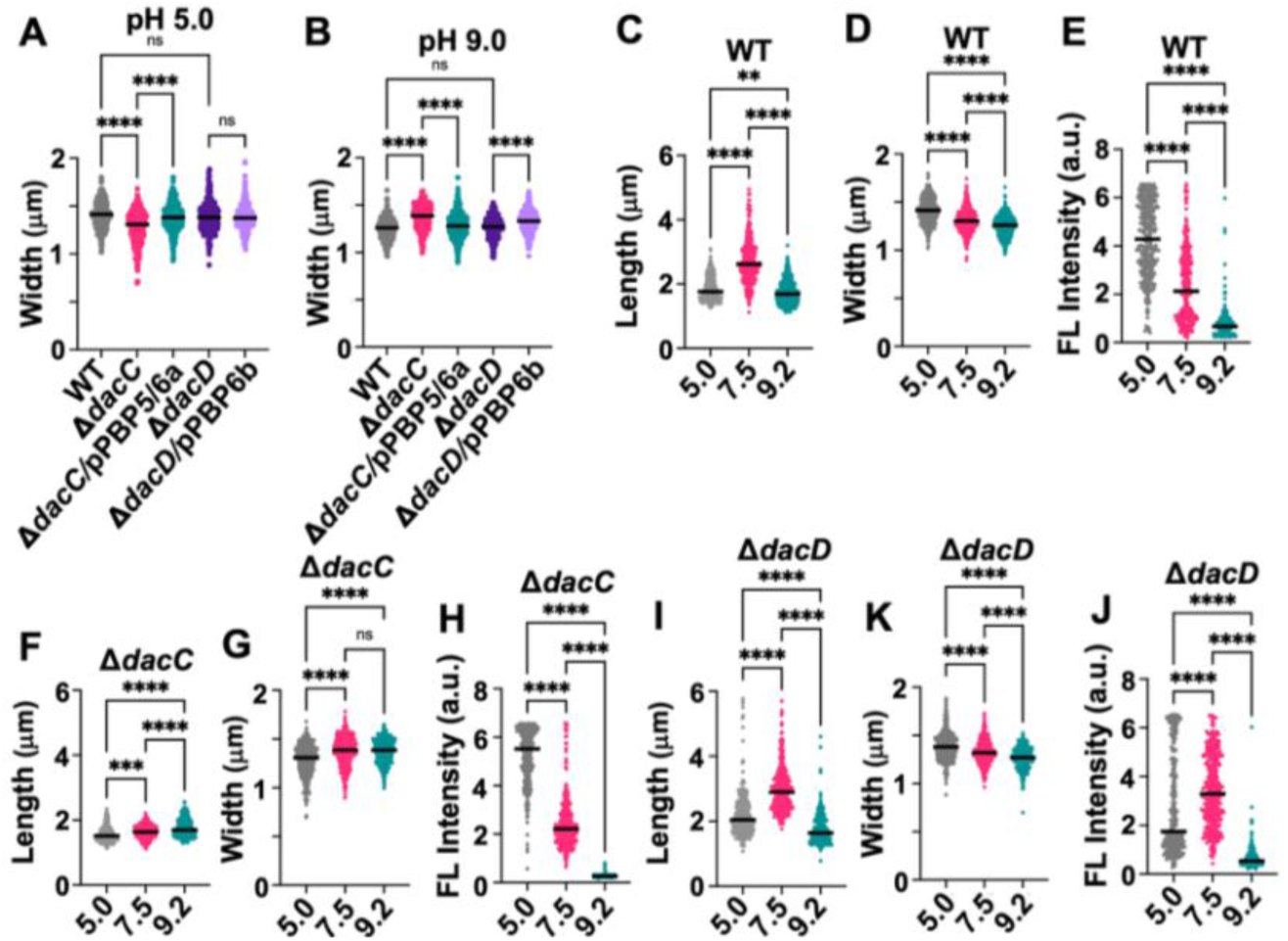
Quantitative analysis of *A. baumannii* strain ATCC 17978 cell morphology and fluorescence under acidic and alkaline conditions. (A) Quantifications of width in wild-type (WT), Δ*dacC*, Δ*dacC*/pPBP5/6a, Δ*dacD*, and Δ*dacD*/pPBP6b strains grown at pH 5.0. **(B)** Same as (A), but for cells grown in pH 9.0. Quantifications (*n* ≥300) were performed using ImageJ with the MicrobeJ plugin. Each dot represents a single cell. Error bars represent standard deviation. Statistical significance was determined using one-way ANOVA (** *P* <0.01, *** *P* <0.001, **** *P* <0.0001). **(C)** Quantification of length, **(D)** width, and **(E)** fluorescence intensity in WT cells grown at pH 5.0, 7.5, and 9.0. **(F, G, H)** Same as described for (C-E), but for Δ*dacC* cells. **(I, J, K)** Same as described for (C-E), but for Δ*dacD* cells.

