## Supplemental Material for "Peptidoglycan DD-peptidases have distinct activities that impact fitness of *Acinetobacter baumannii*"

### Supplemental data for ‘Peptidoglycan DD-peptidases have distinct activities that impact fitness of *Acinetobacter baumannii*’

Arshya Tehrani, Abhisha Khadka, Berenice Furlan, Christine Pybus, Michael Whalen, Jacob Biboy, Orietta Massidda, Waldemar Vollmer, Joseph M. Boll[#]

#### Supplementary Figures

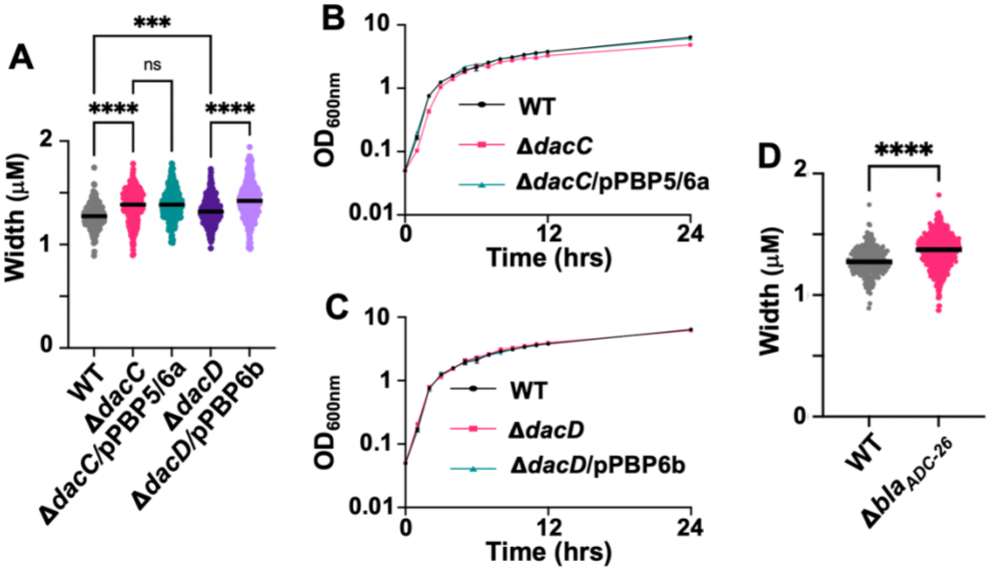


**Figure S1: Analysis of Δ*dacC* and Δ*dacD* mutations in *A. baumannii* strain ATCC 17978.** **(A)** Quantifications of cell width in wild-type (WT), Δ*dacC*, Δ*dacC*/pPBP5/6a, Δ*dacD*, Δ*dacD*/pPBP6b and $\Delta bla_{ADC\text{-}26}$ strains ($n \geq 300$), measured using ImageJ with the MicrobeJ plugin. Each dot represents a single cell. Error bars indicate standard deviation. Statistical significance was determined using one-way ANOVA (*** $P < 0.001$, **** $P < 0.0001$). **(B)** Growth curves of the Δ*dacC* mutant. **(C)** Growth curves of the Δ*dacD* mutant. (**D)** Quantification of cell width in the $\Delta bla_{ADC\text{-}26}$ mutant, as described in (A).

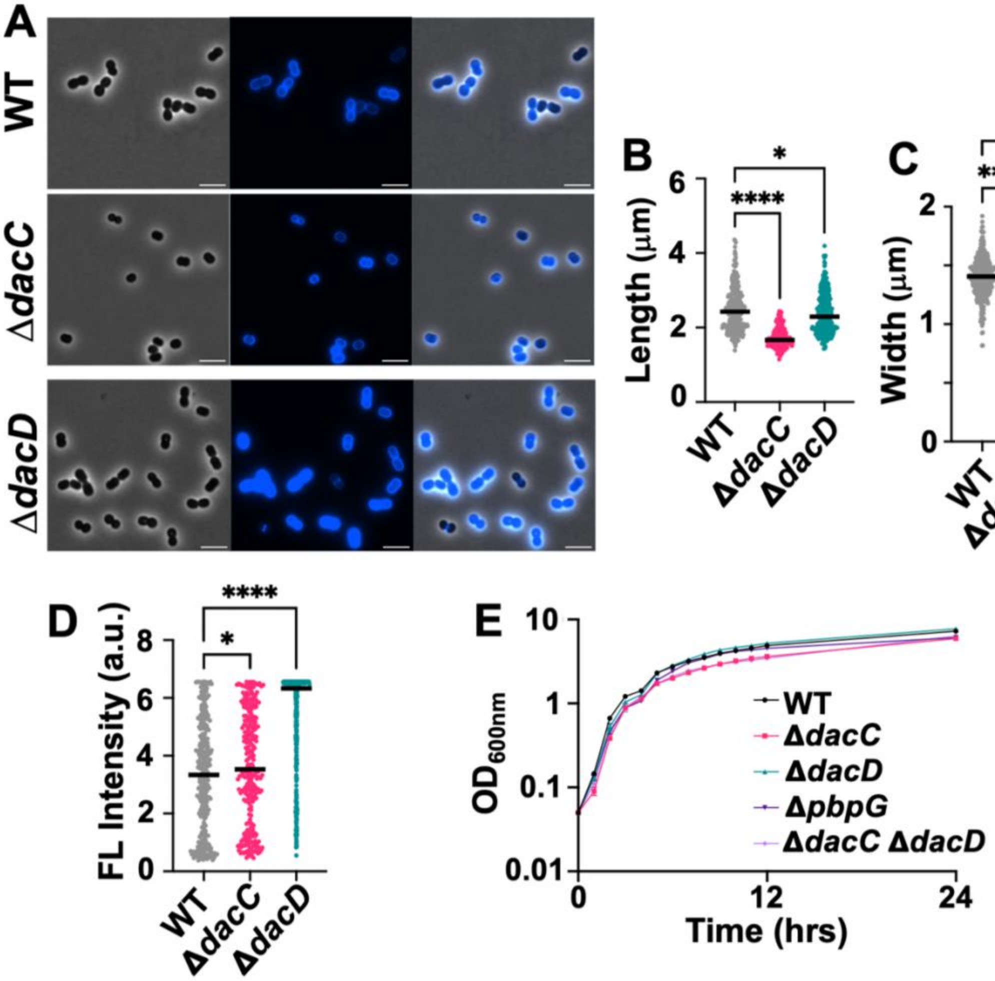


**Figure S2 : Analysis of *A. baumannii* strain AB5075 mutants. (A)** Phase-contrast (left) fluorescence (middle) and merged (right) images of wild-type (WT), Δ*dacC*, Δ*dacD* cells. Scale bar: 5 µm. (**B)** Quantification of cell length (pole to pole) for each strain (*n* ≥300) was measured using ImageJ with the MicrobeJ plugin. Each dot represents a single cell. Error bars indicate standard deviation. Statistical significance was determined using one-way ANOVA ($^{*}$ $P$ < 0.05, $^{****}$ $P$ < 0.0001). **(C)** Quantification of cell width. **(D)** Quantification of fluorescence intensity. **(E)** Growth curves of Δ*dacC,* Δ*dacD,* and Δ*pbpG* mutants. Each experiment was independently replicated three times; one representative data set is shown.

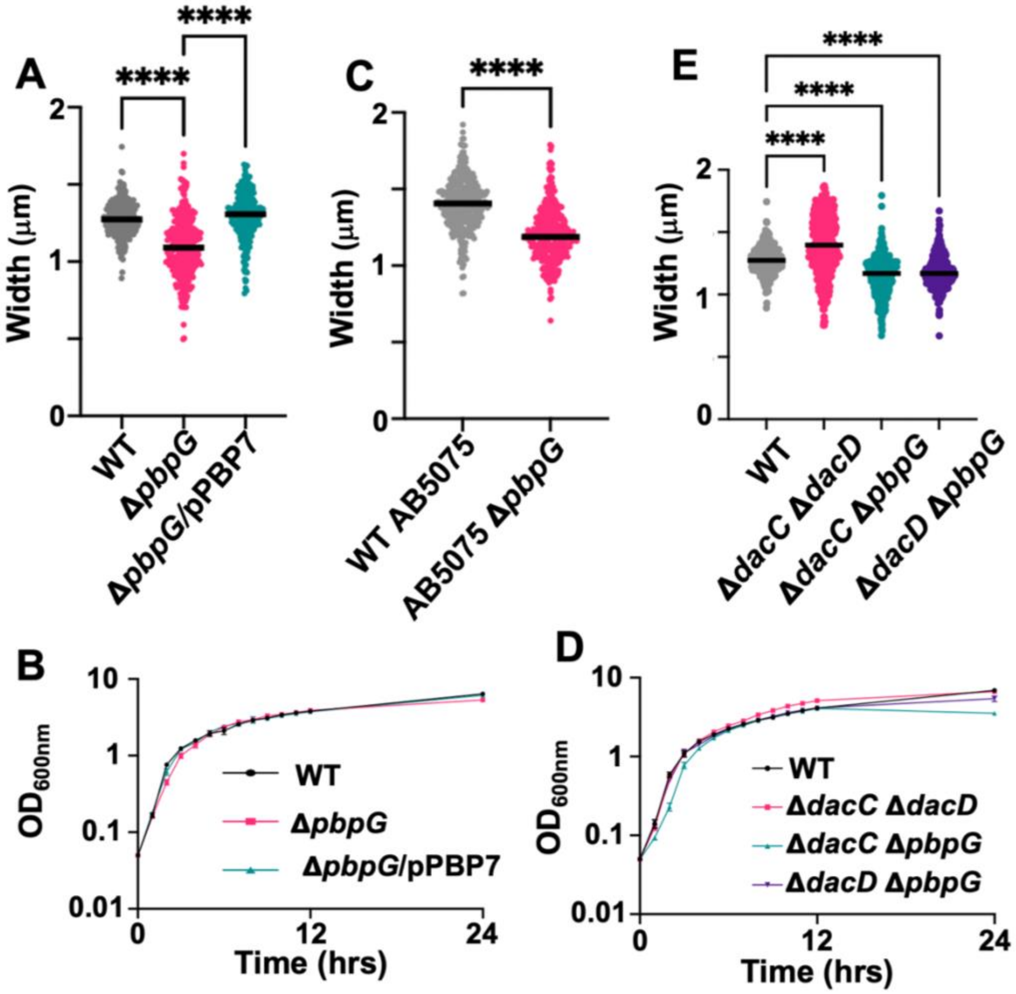


**Figure S3: Cell morphology measurement and growth curves of Δ*pbpG* mutants and double mutants. (A)** Quantifications of cell width in *A. baumannii* strain ATCC 17978 Δ*pbpG* mutants (*n* ≥300) , measured using ImageJ with the MicrobeJ plugin. Each dot represents a single cell. Error bars represent standard deviation. Statistical significance was determined using one-way ANOVA (**** $P < 0.0001$). **(B)** Growth curves of Δ*pbpG* mutants in strain ATCC 17978. **(C)** Same as described in (A), but for Δ*pbpG* mutants in *A. baumannii* strain AB5075 . **(D)** Growth curves of double mutants in strain *A. baumannii* ATCC 17978. **(E)** Same as described in (A), but for double mutants in *A. baumannii* strain ATCC 17978.

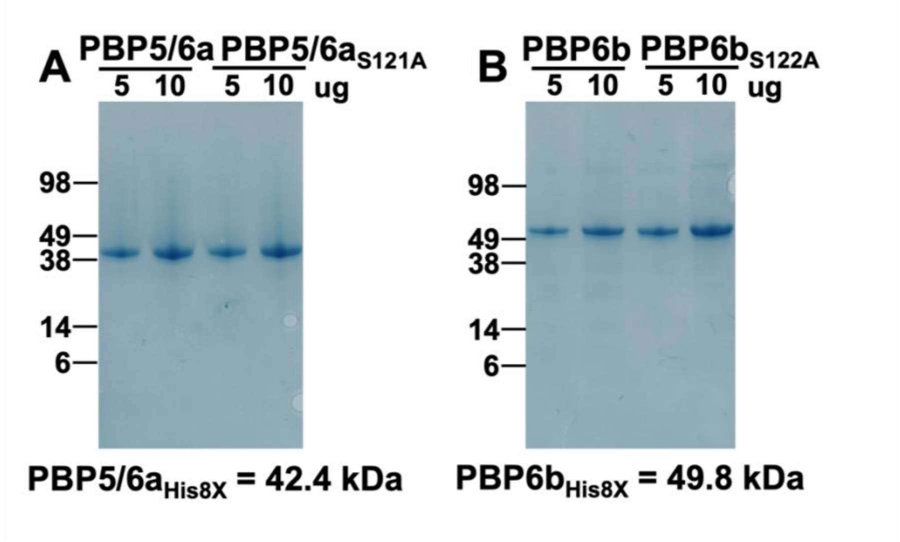


**Figure S4: Purification of PBP5/6a and PBP6b enzymes. (A)** Coomassie stained SDS-PAGE gel of PBP5/6a$_{His8X}$ and the catalytically inactive mutant PBP5/6a$_{S121A\ His8X}$. **(B)** Coomassie stained SDS-PAGE gel of PBP6b$_{His8X}$ and the catalytically inactive mutant PBP6b$_{S122A\ His8X}$.

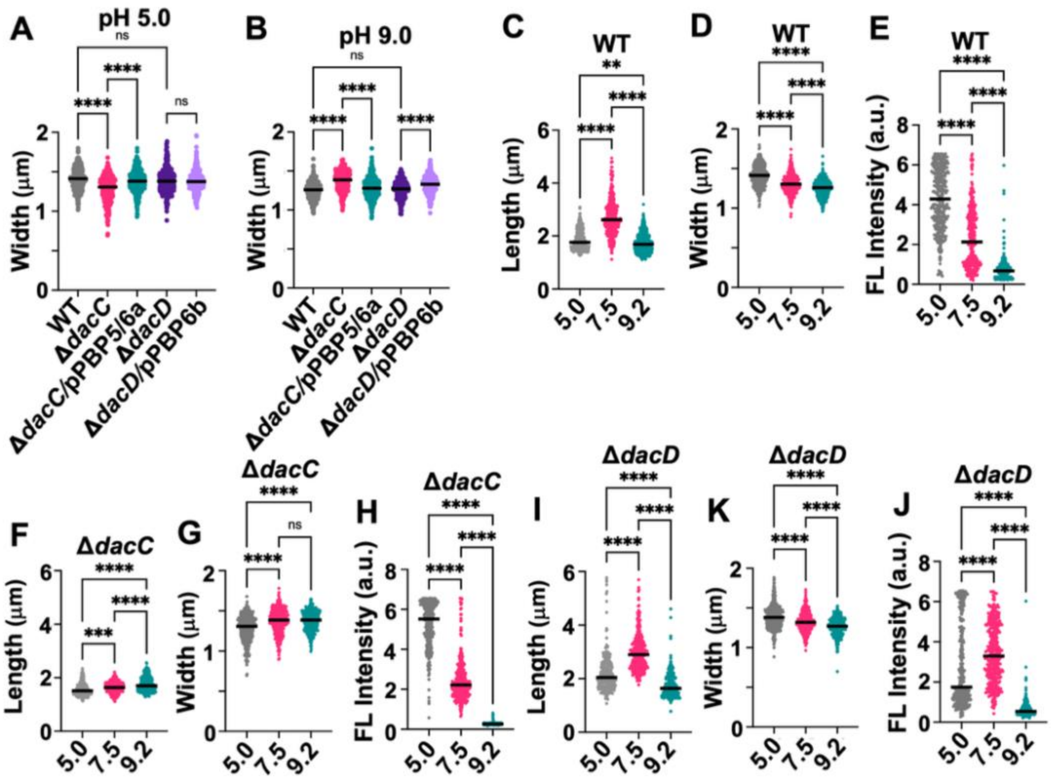


**Figure S5: Quantitative analysis of *A. baumannii* strain ATCC 17978 cell morphology and fluorescence under acidic and alkaline conditions. (A)** Quantifications of width in wild type (WT), Δ*dacC*, Δ*dacC*/pPBP5/6a, Δ*dacD*, and Δ*dacD*/pPBP6b strains grown at pH 5.0. **(B)** Same as (A), but for cells grown in pH 9.0. Quantifications ($n \geq 300$) were performed using ImageJ with the MicrobeJ plugin. Each dot represents a single cell. Error bars represent standard deviation. Statistical significance was determined using one-way ANOVA ($^{**}P$ <0.01, $^{***}P$ <0.001, $^{****}P$ <0.0001). **(C)** Quantification of length, **(D)** width, and **(E)** fluorescence intensity in WT cells grown at pH 5.0, 7.5, and 9.0. **(F, G, H)** Same as described for (C-E), but for Δ*dacC* cells. **(I, J, K)** Same as described for (C-E), but for Δ*dacD* cells.

#### Supplementary Tables

##### Table S1. Established DD-peptidases in *E. coli vs* homologs in *A. baumannii*

| ***E. coli*** | | | | ***A. baumannii*** | | | |
|---|---|---|---|---|---|---|---|
| **Gene** | **Protein** | **Length (aa)** | **Function** | **Gene** | **Protein** | **Length (aa)** | **Predicted function** |
| *dacB* | PBP4 | 477 | DD-CPase; DD-EPase | n/d | n/d | n/d | - |
| *dacA* | PBP5 | 403 | DD-CPase | *dacC* | PBP5/6a | 382 | DD-CPase |
| *dacC* | PBP6a | 400 | DD-CPase | | | | |
| *dac*D | PBP6b | 388 | DD-CPase | *dacD* | PBP6b | 439 | DD-CPase; DD-EPase |
| *pbpG* | PBP7/8 | 479 | DD-EPase | *pbpG* | PBP7 | 348 | DD-CPase; DD-EPase |
| *ampC* | AmpC | 377 | β-lactamase; DD-CPase | $bla_{ADC\text{-}26}$ | AmpC | 383 | β-lactamase; DD-CPase? |
| *ampH* | AmpH/PBP4b | 385 | DD-CPase; DD-EPase | n/d | n/d | n/d | - |

DD-CPase: DD-carboxypeptidase

DD-EPase: DD-endopeptidase

n/d: no homolog detected

**Table S2: Muropeptide composition of wild type (WT), Δ*dacC*, and Δ*dacD* *A. baumannii* strain ATCC 17978**

| Peak No. | Muropeptide | Relative % of each muropeptide[a] | | | | | |
|---|---|---|---|---|---|---|---|
| | | WT Logarithmic | WT Stationary | Δ*dacC* Logarithmic | Δ*dacC* Stationary | Δ*dacD* Logarithmic | Δ*dacD* Stationary |
| 1 | Tri | 2.5 ± 0.0 | 2.9 ± 0.0 | 3.3 ± 0.2 | 4.0 ± 0.1 | 2.6 ± 0.0 | 2.9 ± 0.1 |
| 2 | Tri-D-Asn | 0.7 ± 0.0 | 0.4 ± 0.0 | 0.7 ± 0.0 | 0.6 ± 0.0 | 0.5 ± 0.0 | 1.1 ± 2.2 |
| 3 | Tri-D-Lys | 0.1 ± 0.0 | 0.7 ± 0.1 | 0.5 ± 0.0 | 1.2 ± 1.0 | 0.7 ± 0.0 | 1.5 ± 1.3 |
| 4 | TetraGly4 | 1.8 ± 0.0 | 1.5 ± 0.0 | 2.6 ± 0.0 | 1.3 ± 0.8 | 1.6 ± 0.0 | 1.9 ± 3.1 |
| 5 | Tetra-D-Lys | 0.0 ± 0.0 | 0.2 ± 0.1 | 0.0 ± 0.0 | 0.0 ± 0.0 | 0.4 ± 0.3 | 0.0 ± 0.0 |
| 6 | Tetra | 19.0 ± 0.4 | 20.7 ± 0.0 | 16.9 ± 7.6 | 19.1 ± 0.1 | 16.0 ± 0.4 | 17.6 ± 0.0 |
| 7 | Tetra-D-Arg | 0.0 ± 0.0 | 0.3 ± 0.0 | 0.0 ± 0.0 | 0.4 ± 0.0 | 0.5 ± 0.0 | 1.5 ± 0.0 |
| 7B | Penta | 0.0 ± 0.0 | 0.0 ± 0.0 | 0.5 ± 0.0 | 0.7 ± 0.1 | 0.1 ± 0.0 | 0.0 ± 0.0 |
| 8 | TetraTriDapGly4 | 0.2 ± 0.0 | 0.4 ± 0.0 | 0.5 ± 0.0 | 0.3 ± 0.1 | 0.1 ± 0.0 | 0.1 ± 0.1 |
| 9 | TriTri(Dap)/TriTriDap-D-Lys | 0.5 ± 0.0 | 0.9 ± 0.1 | 0.6 ± 0.0 | 1.3 ± 0.1 | 0.5 ± 0.0 | 0.8 ± 0.2 |
| 10 | TetraTri(Dap) | 0.0 ± 0.0 | 0.2 ± 0.1 | 0.3 ± 0.1 | 0.4 ± 0.0 | 0.2 ± 0.0 | 0.8 ± 1.1 |
| 11 | TetraTri | 3.7 ± 0.1 | 4.0 ± 0.2 | 4.0 ± 0.1 | 4.9 ± 0.6 | 3.1 ± 0.0 | 3.7 ± 0.7 |
| 12 | TetraTri-D-Lys | 0.3 ± 0.1 | 1.9 ± 0.7 | 1.5 ± 0.6 | 2.5 ± 2.0 | 0.9 ± 0.3 | 1.9 ± 1.8 |
| 13 | TetraTri-D-Lys | 0.9 ± 0.1 | 1.0 ± 0.6 | 0.4 ± 0.2 | 1.6 ± 0.2 | 0.4 ± 0.0 | 1.6 ± 0.4 |
| 14 | TetraTri-D-Arg | 0.5 ± 0.0 | 1.0 ± 0.4 | 1.4 ± 0.4 | 1.1 ± 1.3 | 2.1 ± 0.1 | 3.4 ± 19.2 |
| 15 | TetraTetra | 37.7 ± 1.4 | 31.5 ± 1.0 | 32.5 ± 5.8 | 25.8 ± 0.8 | 34.7 ± 0.6 | 25.4 ± 24.5 |
| 15B | TetraPenta | 0.4 ± 0.0 | 0.7 ± 0.1 | 1.6 ± 0.0 | 1.8 ± 0.0 | 1.6 ± 0.4 | 2.8 ± 0.0 |
| 16 | TetraTetraTri or TetraTetraTriDap | 1.1 ± 0.0 | 1.7 ± 0.0 | 1.0 ± 0.5 | 2.0 ± 0.0 | 1.1 ± 0.0 | 1.4 ± 0.0 |
| 17 | TetraTetraTri or TetraTetraTriDap | 0.2 ± 0.0 | 0.6 ± 0.1 | 0.8 ± 0.2 | 1.7 ± 1.0 | 0.6 ± 0.1 | 0.6 ± 0.1 |
| 18 | TriTriDap-D-Met | 0.8 ± 0.0 | 1.1 ± 0.2 | 1.0 ± 0.8 | 0.7 ± 0.2 | 0.3 ± 0.2 | 1.1 ± 0.1 |
| 19 | TetraTetraTetra | 18.5 ± 0.0 | 16.1 ± 0.1 | 15.2 ± 7.2 | 12.8 ± 0.0 | 17.8 ± 0.0 | 13.6 ± 0.1 |
| 20 | TetraTri-D-Met | 0.4 ± 0.0 | 0.5 ± 0.0 | 0.6 ± 0.0 | 1.0 ± 0.1 | 0.3 ± 0.0 | 0.5 ± 0.1 |
| 21 | TetraTriAnh / TetraTetraTetraTri | 4.8 ± 0.0 | 4.2 ± 0.0 | 3.7 ± 1.8 | 3.3 ± 0.0 | 4.7 ± 0.0 | 3.5 ± 0.1 |
| 22 | TetraTetraAnh I | 1.7 ± 0.2 | 1.3 ± 0.1 | 1.8 ± 0.8 | 1.1 ± 0.2 | 1.6 ± 0.3 | 1.2 ± 0.2 |
| 23 | TetraTetraAnh II | 0.7 ± 0.0 | 0.9 ± 0.0 | 0.8 ± 0.0 | 0.9 ± 0.0 | 0.7 ± 0.0 | 0.9 ± 0.1 |
| 24 | TetraTetraTetraAnh | 2.0 ± 0.0 | 2.1 ± 0.0 | 2.0 ± 0.0 | 1.7 ± 0.1 | 2.1 ± 0.0 | 2.0 ± 0.0 |
| Sum of known peaks | | 97.7 ± 1.1 | 96.0 ± 3.9 | 93.2 ± 27.4 | 91.5 ± 0.0 | 94.7 ± 0.7 | 90.9 ± 0.3 |
| Monomers (Total) | | 24.5 ± 0.7 | 27.7 ± 0.0 | 26.2 ± 19.8 | 29.6 ± 0.0 | 23.5 ± 0.0 | 28.9 ± 5.4 |
| Monomers with modification | | 2.6 ± 0.0 | 3.2 ± 0.0 | 4.0 ± 0.0 | 3.7 ± 0.2 | 3.8 ± 0.1 | 6.4 ± 4.5 |
| Monomer tri | | 2.5 ± 0.0 | 3.0 ± 0.0 | 3.5 ± 0.5 | 4.4 ± 0.1 | 2.7 ± 0.1 | 3.2 ± 0.1 |
| Monomer tri-D-Asn | | 0.7 ± 0.0 | 0.4 ± 0.0 | 0.8 ± 0.0 | 0.6 ± 0.0 | 0.5 ± 0.0 | 1.1 ± 2.4 |
| Monomer tri-D-Lys | | 0.1 ± 0.0 | 0.7 ± 0.1 | 0.5 ± 0.0 | 1.3 ± 1.3 | 0.7 ± 0.0 | 1.7 ± 1.6 |
| Monomer tetraGly4 | | 1.9 ± 0.0 | 1.5 ± 0.0 | 2.7 ± 0.0 | 1.4 ± 1.0 | 1.7 ± 0.0 | 2.1 ± 3.6 |
| Monomer tetra-D-Lys | | 0.0 ± 0.0 | 0.3 ± 0.1 | 0.0 ± 0.0 | 0.0 ± 0.0 | 0.4 ± 0.3 | 0.0 ± 0.0 |
| Monomer tetra | | 19.4 ± 0.7 | 21.5 ± 0.1 | 18.2 ± 15.7 | 20.9 ± 0.2 | 16.9 ± 0.2 | 19.3 ± 0.0 |
| Monomer tetra-D-Arg | | 0.0 ± 0.0 | 0.3 ± 0.0 | 0.0 ± 0.0 | 0.4 ± 0.0 | 0.6 ± 0.0 | 1.6 ± 0.0 |
| Monomer penta | | 0.0 ± 0.0 | 0.0 ± 0.0 | 0.5 ± 0.0 | 0.8 ± 0.1 | 0.1 ± 0.0 | 0.0 ± 0.0 |
| Dimers (Total) | | 53.2 ± 1.0 | 51.3 ± 0.2 | 53.7 ± 6.1 | 50.7 ± 0.6 | 53.9 ± 0.0 | 52.0 ± 5.4 |
| Dimers with modification | | 3.4 ± 0.2 | 6.9 ± 0.0 | 6.0 ± 3.4 | 9.1 ± 0.2 | 4.9 ± 0.1 | 10.1 ± 8.4 |
| Dimer chain ends (anhydroMur*N*Ac) | | 7.3 ± 0.5 | 6.5 ± 0.1 | 6.6 ± 0.0 | 5.7 ± 0.2 | 7.4 ± 0.4 | 6.1 ± 0.2 |
| Trimers (Total) | | 22.3 ± 0.0 | 21.2 ± 0.0 | 20.3 ± 4.2 | 19.9 ± 0.6 | 22.8 ± 0.0 | 19.2 ± 0.0 |
| Trimer chain ends (anhydroMur*N*Ac) | | 2.1 ± 0.0 | 2.1 ± 0.0 | 2.2 ± 0.0 | 1.9 ± 0.1 | 2.3 ± 0.0 | 2.2 ± 0.0 |
| Tripeptides (Total) | | 10.0 ± 0.1 | 13.0 ± 0.3 | 12.9 ± 1.8 | 17.0 ± 4.5 | 11.2 ± 0.1 | 16.5 ± 3.4 |
| Tripeptides with modifications | | 3.1 ± 0.1 | 5.6 ± 0.4 | 5.1 ± 2.2 | 7.5 ± 1.0 | 4.1 ± 0.0 | 8.8 ± 1.6 |
| Tetrapeptides (Total) | | 89.5 ± 0.1 | 86.0 ± 0.7 | 85.3 ± 2.4 | 80.5 ± 3.9 | 87.6 ± 0.0 | 81.5 ± 2.4 |
| Tetrapeptides with modifications | | 2.9 ± 0.1 | 4.5 ± 0.4 | 4.9 ± 0.2 | 5.3 ± 1.0 | 4.7 ± 0.2 | 7.7 ± 14.0 |
| Pentapeptides | | 0.2 ± 0.0 | 0.3 ± 0.0 | 1.3 ± 0.0 | 1.7 ± 0.0 | 1.0 ± 0.0 | 1.5 ± 0.0 |
| 3-3 Crosslinks | | 1.1 ± 0.0 | 2.1 ± 0.0 | 1.8 ± 0.6 | 2.8 ± 0.0 | 1.2 ± 0.1 | 2.2 ± 0.1 |
| Chain ends (anhydroMur*N*Ac) | | 4.3 ± 0.1 | 4.0 ± 0.0 | 4.0 ± 0.0 | 3.5 ± 0.1 | 4.4 ± 0.1 | 3.8 ± 0.0 |
| Degree of crosslinkage | | 41.4 ± 0.2 | 39.7 ± 0.0 | 40.3 ± 6.5 | 38.6 ± 0.0 | 42.1 ± 0.0 | 38.8 ± 1.4 |
| % peptides in cross-links | | 75.5 ± 0.7 | 72.4 ± 0.0 | 73.9 ± 19.8 | 70.5 ± 0.0 | 76.6 ± 0.0 | 71.2 ± 5.4 |

[a]Values are mean ± variation of two biological repeats.

**Table S3: Strains and plasmids used in this study**.

| Strain/Plasmid | Description | Reference/Source |
|---|---|---|
| **Strains** | | |
| *E. coli* C2987 | chemically competent wild type, K-12 | New England Biolabs |
| *E. coli* C2527 | chemically competent BL-21 | New England Biolabs |
| *A. baumannii* ATCC 17978 | wild type | ATCC (1) |
| *A. baumannii* 5075 | wild type | (2) |
| *A. baumannii* ATCC 17978 | Δ*dacC* | This Study |
| *A. baumannii* ATCC 17978 | Δ*dacC*/pPBP5/6a | This Study |
| *A. baumannii* ATCC 17978 | Δ*dacD* | This Study |
| *A. baumannii* ATCC 17978 | Δ*dacD*/pPBP6b | This Study |
| *A. baumannii* ATCC 17978 | Δ*pbpG* | (3) |
| *A. baumannii* ATCC 17978 | Δ*pbpG*/pPBP7 | (3) |
| *A. baumannii* ATCC 17978 | Δ$bla_{ADC\text{-}26}$ | This Study |
| *A. baumannii* ATCC 17978 | Δ*dacC* Δ*dacD* | This Study |
| *A. baumannii* ATCC 17978 | Δ*dacC* Δ*pbpG* | This Study |
| *A. baumannii* ATCC 17978 | Δ*dacD* Δ*pbpG* | This Study |
| *A. baumannii* 5075 | Δ*dacC* Δ*dacD* | This Study |
| *A. baumannii* 5075 | Δ*dacC* Δ*pbpG* | This Study |
| *A. baumannii* 5075 | Δ*dacD* Δ*pbpG* | This Study |
| **Plasmids** | | |
| pABBRKn | pABBR_MCS with the $Kan^R$ gene from pKD4 inserted into the PvuI site, $Kn^R$ | (5) |
| pAT03 | pMMB67EH with FLP recombinase, $Amp^R$ | (4) |
| pAT04 | pMMB67EH with $REC_{Ab}$ system, $Tet^R$ | (4) |
| pKD4 | $Kan^R$ | (6) |
| pT7-7 | $Amp^R$ | (7) |
| pT7-7Kn | pT7-7 with the $Kan^R$ gene from pKD4 inserted into the PvuI site, $Kn^R$ | (3) |
| pPBP5/6a-$His_{8X}$ | pT7-7 with *dacC* (A1S_2435) cloned into the NdeI and BamHI sites, $Kn^R$ | This study |
| pPBP5/6$a_{S121A}$-$His_{8X}$ | pT7-7 with $dacC_{S121A}$ cloned into the NdeI and BamHI sites, $Kn^R$ | This study |

| | | |
|---|---|---|
| pPBP7 | pMMB67EHKn with the *pbpG* (A1S_0237) gene and native promoter inserted into the XhoI and KpnI sites, Kn$^R$ | (3) |
| pPBP6b-His$_{8X}$ | pT7-7 with *dacD* (A1S_2479) cloned into the NdeI and BamHI sites, Kn$^R$ | This study |
| pPBP6b$_{S122A}$-His$_{8X}$ | pT7-7 with *dacD*$_{S122A}$ cloned into the NdeI and BamHI sites, Kn$^R$ | This study |
| pPBP5/6a | (A1S_2435) gene and native promoter inserted into the XhoI and KpnI sites, Kn$^R$ | This study |
| pPBP6b | (A1S_2479) gene and native promoter inserted into the XhoI and KpnI sites, Kn$^R$ | This study |

**Table S4: Oligonucleotides used in this study.**

| **Oligo Name** | **Sequence (5' to 3')** |
|---|---|
| **Deletion Primers** | |
| 17978 *dacC* Kan-FRT 5' | GTTAAAAACTCTCTAGTTACAATACTGTTTGAAAAAGCCGACTTCCCCCATAGAAGTCGGCTTTGCTTTATCCTAAAATGCTAGGCTCTTCTGCTTCATACAAGATATTGGAATTACCTAGAATGATATCCTCCTTAGTTCCTATTCCG |
| 17978 *dacC* Kan-FRT 3' | AACTTTACCATCTAAACTTGCAACAAGCTTACCAACGACTTGACCTTTTTGAAGTGGTGCATTTAGGTTCGGTTGAACAACCAATTGAGTTTTAATGCCGTCCGCTTTGCCTTTAGGCATAGTTAAGCGATTGTGTAGGCTGGAGCTGCTTCG |
| 17978 *dacC*-kan verify 5' | AGCACGTAATGATGCAAAAGCTG |
| 17978 *dacC*-kan verify 3' | CTTGGATCAATTTGTGGGTGATGGTC |
| 17978 *dacD* Kan-FRT 5' | TTTGTATCGACTTGCGTAGATGTATTTTTTGGCAACAATTCACACTAGCCTTCTACAAAAAAAAGGCATAGCATGCTGGGCATCTGTGTTCCGTATTTGCCAGTAAACGTCAAGGTCACCTAGTG |
| 17978 *dacD* Kan-FRT 3' | TTTGCTTCTTCAATGTGTACATCATTTTCAATTTGAAGACTGCGAATGAGCTGGTTGTTTTGATAAATTGAAACTGTTGCCAAATTCATCGCTTTCATTAACGGTGCTGTTAACTTTTGCTCATT |
| 17978 *dacD*-kan verify 5' | GCGAATTGTCACGTGAACAAGG |
| 17978 *dacD*-kan verify 3' | GCGTGGCGAGTTCTAAACCAC |
| 17978 *pbpG*-Kan FRT 5' | AAAGCTTTATACCTTATATCTCAAATGTAAGGCATAATGATAGTAAGCGCAAATGTGTGTCACCCTGAGTCGAGTATTGTGCCGTGAAAAATTCTAAAAAGTCTTTAATGCATGTGCTAAGCATGatatcctccttagttcctattccg |
| 17978 *pbpG* -Kan FRT 3' | TATCTAATATGTAAAATCTGAGTTTTTATAAAAAGC |

| | |
|---|---|
| | GGCTGTTTAATACAGCCGTTTTTTTATGCTTTTTAAATGGCATAAAAAAACGCTTCTTAAAAGAAGCGTTTTTAAAAATAATTAAATTAagcgattgtgtaggctggagctgcttcg |
| 17978 *pbpG*-kan verify 5’ | GCTTGCAATGGAATGACAAAATTAGCAATC |
| 17978 *pbpG*-kan verify 3’ | GATACAGCAATTAAACAATGTGCTGATGCAG |
| 17978 $bla_{ADC\text{-}26}$ Kan-FRT 5’ | GCTTTTTTTATAGCTGAACGCGATAAACTTCAATCCAATTCTATAAAACATAACAATTAAATTAGGTGATTTTTGTTATAAAAGTAGGCATCTTTCTTTTTAAATAATTTATGAGCTAATCagcgattgtgtaggctggagctgcttcg |
| 17978 $bla_{ADC\text{-}26}$ Kan-FRT 3’ | CAAAATCATCTGGACTGGATGAAATCAAGCTTAGGATATGTTTGGTTCTTTTAAACCATATACCTATAAAAAATAGAGAAATAGATGATGACTATTTCTCTCTTTTTATTTTAGGGAAAAAatatcctccttagttcctattccg |
| 17978 $bla_{ADC\text{-}26}$-kan verify 5’ | GTGTATGATGATGCGTGGTGTAGGT |
| 17978 $bla_{ADC\text{-}26}$-kan verify 3’ | GTGTGGGAGTAGGGCCGAC |
| **Complementation Primers** | |
| pABBR-*dacC*-F XhoI | CGCCTCGAGatgcaaaagctgctggtaacgaac |
| pABBR-*dacC*-R KpnI | CGCGGTACCccacggacatttcatcaatgtaacagc |
| pABBR-*dacD*-F XhoI | CGCCTCGAGttcctgatttggattcgctaaaaagcg |
| pABBR-*dacD*-R KpnI | CGCGGTACCggctttaccatttttcccaaaagtttcacc |
| pABBR-*pbpG*-F XhoI | AAATTACTCGAGCTATTCTTCTATAGTGAGCGAATAGTTG |
| pABBR-*pbpG*-R KpnI | GTTGTCGGTACCTGCAACAATGGACCAAGTAAAAGATTCG |
| **Overexpression Primers** | |
| pT7-*dacC* NdeI | CGCCATATGACTCGAAAAAGCGCTATTGCTGCACTCCTCCTCTTAC |
| pT7-*dacC* BamHI 8X-his | CGCGGATCCTTAATGGTGATGGTGATGGTGATGGTGGTTGCTGAAGAATTGTTTGATATGG |
| pT7-*dacD* NdeI | CGCCATATGAAATTCTTCCTATCTCTTTTTACGCTGTTTAGTATTTTCTGTACTACTCTTACC |
| pT7-*dacD* BamHI 8X-his | CGCGGATCCTTAATGGTGATGGTGATGGTGATGGTGGTGCGAATCTATAGGG |
| **Sequencing Primers** | |
| pABBR confirm 1 | GGGCTGACCGCTTCCT |
| pABBR confirm 2 | CGCTAGCAGCACGCCATAG |
